# High activity and high functional connectivity are mutually exclusive in resting state zebrafish and human brains

**DOI:** 10.1101/2021.06.02.446741

**Authors:** Mahdi Zarei, Dan Xie, Fei Jiang, Adil Abigrove, Bo Huang, Ashish Raj, Srikantan Nagarajan, Su Guo

## Abstract

Active neurons impact cell types with which they are functionally connected. Both activity and functional connectivity are heterogeneous across the brain, but the nature of their relationship is not known. Here we employ brain-wide calcium imaging at cellular resolution in larval zebrafish to record spontaneous activity of >12,000 neurons in the forebrain. By classifying their activity and functional connectivity into three levels (high, medium, low), we find that highly active neurons have low functional connections and highly connected neurons are of low activity. Intriguingly, deploying the same analytical methods on functional magnetic resonance imaging (fMRI) data from the resting state human brain, we uncover a similar relationship between activity and functional connectivity, that is, regions of high activity are non-overlapping with those of high connectivity. These findings reveal a previously unknown and evolutionarily conserved brain organizational principle that have implications for understanding disease states and designing artificial neuronal networks.

## Introduction

The structure of the brain spans dimensions that are many orders of magnitudes apart, from molecules, synapses, cells, to meso- and macro-scale brain systems. Such extraordinary architecture serves not only to process sensory information that guides motor behaviors, but also to generate and maintain internal states (e.g., emotional, motivational, and cognitive states) that can critically influence an organism’s response to environment.

Diverse approaches have been employed to understand the brain’s architecture at both anatomical and functional levels across multiple scales in invertebrate and vertebrate organisms (*1–7*). Viral tracing and MRI/fMRI studies provide meso-to macro- scale descriptions of connectivity in mammalian brains (*8–10*). A single-cell resolution connectome of *C. elegans*’ 302 neurons is constructed at the nanometer scale using electron microscopy (EM) (*1*), but the application of EM to more complex brains requires enormous time and resources, making it best suited for small parts of brain tissues from one or few individuals (*11–14*). While structural connectome is foundational to functional connectome, it is the functional connectome that underlies behavioral and mental states. One effective way to gain insights into brain’s functional architecture is to record spontaneous neuronal activities brain-wide and analyze their activity and functional connectivity. Functional connectivity measures correlations between time series of individual neurophysiological events (*15*). Such connectivity may be direct, indirect through a subnetwork (*16*), or via wireless neuro-modulatory communications (*17*). Brain-wide functional connectivity studies have been mainly carried out in humans using blood-oxygen-level-dependent (BOLD) fMRI (*7*) and MEG/EEG data (*18*). Recent technological advancements in neural activity reporters (*19*) and fast *in vivo* imaging technologies (*20–22*) have made it possible to record whole brain activity at cellular resolution in larval zebrafish (*3, 23, 24*), a vertebrate model organism with relatively small and transparent brains. An elegant body of work has uncovered brain-wide dynamics underlying sensorimotor behaviors (*3, 25–30*). Studies of spontaneous neuronal activities in zebrafish however have been few. Nevertheless, these studies, mostly focused on the larval optic tectum, have revealed that spontaneous activity represents “preferred” network states with propagating neuronal avalanches (*31, 32*). Spontaneous activity can be reorganized over development (*33*), and reflects a spatial structure independent of and activate-able by visual inputs (*34*).

The dynamicity of activity and functional connectivity patterns in the resting state brain have long fascinated systems neuroscientists (*35*). The resting state brain activity refers to spontaneous activity without deliberately given stimuli. Such activity shows relatively consistent distributed patterns and can be used to characterize network dynamics without needing an explicit task to drive brain activity. Analyses of cross-correlation between activity in different brain regions demonstrate that resting state networks (RSNs) (*36*) and default mode networks (DMNs) (*37*) reflect to a considerable extent the anatomical connectivity between the regions in a network. Such intrinsic activity dynamics is shown to be disrupted in neuropsychiatric disorders (*38*).

In this study, we exploit the resting-state brain activity to ask a previously unaddressed question: how does the activity of a neuron (or neuronal population) predict the extent of its functional connectivity? Since neuronal activity is an essential drive that underlies functional connectivity, we hypothesize that neurons with high activity will likely have high functional connectivity. To test this hypothesis, we applied selective-plane illumination microscopy (SPIM) (*20, 39*) to image individual neuron’s spontaneous activity across the forebrain of transgenic larval zebrafish expressing nuclear targeted GCAMP6s. The vertebrate forebrain shares considerable homology in developmental ontogeny and gene expression domains, and is functionally involved in sensory, emotional and cognitive processing (*40–43*). In 6-day old larval zebrafish, the forebrain is spontaneously active with strong local correlations and relatively reduced long-range correlations with the mid- and hindbrain areas (*24*),(*27*). It remains unclear how such spontaneous activity in the forebrain informs the underlying functional architecture. Employing image processing methods to detect individual neurons, we obtained time-dependent activity data for more than 12K neurons per individual forebrain. Through image registration to a brain atlas (*44*), we assigned anatomical labels to each neuron. We established methods to identify optimal thresholding values, at which the functional connectivity was computed. By further classifying individual neurons into three activity and connectivity groups (high, medium, and low), we uncovered a surprising complementary distribution of highly active vs. highly connected neurons. Moreover, we extended such analytical methods to fMRI datasets from the resting state human brain and results like that of zebrafish were obtained. Together, these findings reveal a mutually exclusive relationship between high activity and high functional connectivity. Its plausible cause and implications are discussed in the Discussion section below.

## Results

### Light-sheet imaging and image processing generate large-scale single neuron activity data across the larval zebrafish forebrain

Using a light-sheet imaging system custom constructed based on the iSPIM design (*39*), we recorded the spontaneous activity of neurons in the larval zebrafish forebrain under awake resting state. For each individual, calcium imaging data were collected at ~ 2 volumes per second (26 Z planes with 4 μm interval per volume) for ~15 min (n=9), and acquired images were processed via an image processing pipeline, resulting in a set of multi-dimensional data (Figure S1A-B, Videos S1). The pan-neuronally expressed calcium indicator GCAMP6s fused to the histone H2B protein was localized to the cell nuclei. Using this feature, we segmented the brain into individual neurons (Figure S1C). Neuronal activity as reflected by ΔF/F was calculated using time-dependent baseline estimation procedure as previously described (*45*).

In any calcium imaging experiment, fluorescent signal changes as a measure of neuronal activation are often plagued with noises, either from the instrument or from baseline fluctuations. To differentiate genuine neuronal activity-related peaks from such background noises, we applied a method based on the Bayesian inference of two-dimensional distribution of adjacent ΔF/F values (*31*). This enabled us to obtain “ultra-cleaned” data at 99% confidence levels (Figure S1D). Together, these experiments generate large-scale single neuron activity data across a healthy group of individuals at the awake resting state (Figure S1E).

### Brain registration enables comparison of anatomically identifiable neuronal activity patterns across different individual larval zebrafish

Since individuals differ in morphology, position orientation under the imaging microscope and GCAMP signal intensity, it is difficult to directly compare their brain activity data even though such data are acquired under identical conditions to the experimenter’s knowledge. In order to compare data across individuals, we registered the imaging stacks to the Z-brain atlas (*44*). The iSPIM imaging stacks, which are acquired at a 45-degree angle to the anteroposterior axis, however, cannot be directly registered to the Z-brain template, due to: 1) a significant mismatch between the image directions of our stacks and the Z-brain template, and 2) a significant difference between the volumes of interest (our highly sampled forebrain vs. the whole brain). To address this problem, we created an intermediate reference brain from the Z-brain template, by resampling the forebrain region in the direction and pixel sizes that are comparable to those of the iSPIM stacks (Figure S2A-B). For each individual, a densely sampled Z-stack (with 1 μm interval) was collected and used for registration to the intermediate reference brain using computational morphometry toolkit (CMTK)(http://nitrc.org/projects/cmtk) (*46*). An example of pre- and post-registration images were shown in Figure S2C.

We assigned the 294 anatomical masks in the Z-brain template to the registered iSPIM stacks by reformatting the coordinates for each detected neuron according to the registered frame. This process enabled us to identify anatomical labels for each neuron and the brain regions covered by our imaging volumes (Figure S3). Taken together, these analyses generate anatomically identifiable neuronal activity data that can be compared across individuals.

### Visualization of neuronal activity landscape at single-cell resolution in the larval zebrafish forebrain

As a first step toward data analysis, we visualized the neuronal activity landscape (Fig. 1A). Each neuron’s level of activity was calculated based on the variances of ΔF/F across time. The k-means clustering, which is a well-known unsupervised learning algorithm (*47*), was used to group neurons based on their activity levels. To distinguish neurons with the highest or lowest levels of activity, we set the number of clusters to 3. Hence, the activity levels of 1, 2, and 3 denoted neuronal groups with high, medium, and low activity. The activity level before and after classification was shown for an example subject (Fig. 1B). More than 80% of neurons were classified as Activity Level 3 (AL-3), whereas only ~2% of neurons belonged to Activity Level 1 (AL-1) (Fig. 1C, 1E). Visualization of their anatomical distribution showed that AL-1 neurons were mostly located in the lateral region of the forebrain, whereas AL-3 neurons were distributed in all brain areas (Fig. 1D). Similar observations were made across all subjects, as reflected by the population statistics (Fig. 1F) and the overlay view of AL-1 neurons from all subjects (Fig. 1G). Analysis of detailed anatomical distributions for AL-1 neurons showed that they are mostly located in the telencephalic pallium and diencephalic habenula (Fig. 1H). Together, these findings uncover highly active neurons that are located laterally in the larval zebrafish forebrain.

**Figure 1.**
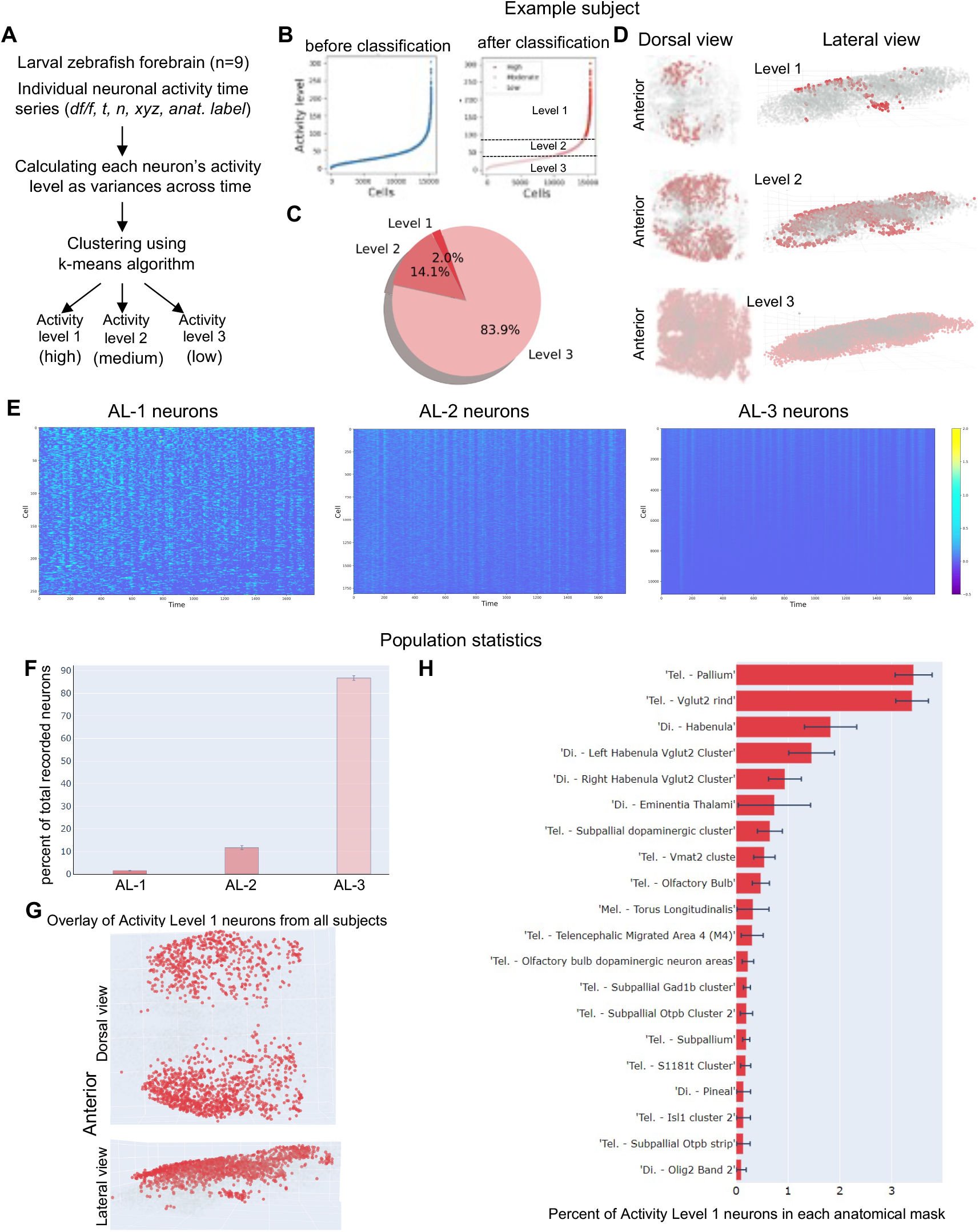
Visualization of neuronal activity landscape at cellular resolution in the larval zebrafish forebrain. **A,** Overview of the classification of individual ROIs (neurons) based on their level of activity. The variance of df/f of each ROI was used as a measure of its activity. The k-means algorithm was used to classify each ROI into 3 levels. **B,** Sorted ROIs (left) vs. clustered ROIs (right) based on their activity level for an example subject. **C,** Pie chart showing the percentage of ROIs in three activity level categories for an example subject. Less than 2% of ROIs are highly active (level I) but more than 80% are largely inactive. **D,** Dorsal and lateral views of the three activity categories of ROIs’ distributions in the example subject's forebrain. **E,** Raster plot of ROIs with different levels of activity in an example subject: (left) Activity Level 1; (right) Activity Level 3. **F,** Percent of total recorded neurons in each activity level category across 9 subjects. **G,** overlay view of highly active neurons (Level 1) in all 9 subjects shows that they are located in the lateral part of the forebrain. **H**, Anatomical distribution of Activity Level 1 neurons sorted based on the percentage of total recorded neurons in each anatomical mask.

### Classification of neurons based on their levels of functional connections in the larval zebrafish forebrain

The brain as a complex network involves intricate communications between individual neurons. An understanding of their patterns of communications will likely inform underlying network architectures. We therefore classified neurons based on their levels of functional connections. Here the degree or number of connections between a neuron and the rest of neurons in the dataset is used as a measure of functional connectivity, which can be approximated using various statistical measures. One common and effective measure for estimation of the connectivity matrix is the Pearson correlation coefficient value. We calculated the degree of connections for each neuron by applying optimal thresholding to the connectivity matrixes followed by binarization. The optimal thresholding value for each subject was determined using the principle of small world networks that follow a power law distribution (*48, 49*) (Fig. S4). Such power law distribution was not observable in randomly shuffled data (Fig. S5), indicating its biological relevance. Moreover, we showed that known connections between olfactory epithelial and olfactory bulb neurons were uncovered (Fig. S6), thereby validating our method of detecting functional connections.

We next used the k-means algorithm to cluster neurons based on their degree of functional connections (Fig. 2A), with the number of clusters also set to 3. The connectivity level before and after k-means classification was shown for an example subject (Fig. 2B). ~8% of neurons belonged to Connectivity Level 1 (CL-1, the high connectivity group) whereas ~70% of neurons were classified as Connectivity Level 3 (CL-3, low connectivity)(Fig. 2C). Visualization of their anatomical distributions showed that the CL-1 neurons were mostly located in the medial area of the forebrain (Fig. 2C). Two example CL-1 neurons had 1814 and 1506 functional connections respectively, in contrast to two example CL-3 neurons with 10 and 19 connections respectively (Fig. 2E). Consistent with the example subject, population statistics showed that the CL-1 neurons represent ~8% of total recorded forebrain neurons (Fig. 2F) and they are located in the medial region of the forebrain (Fig. 2G). Analysis of detailed anatomical distributions for CL-1 neurons uncovered that Telencephalic Olig2 Cluster, Telencephalic S1181t Cluster, and Telencephalic subpallial Otpb strip are among the neuronal groups with high degrees of functional connections in the zebrafish forebrain (Fig. 2H). Together, these findings uncover neurons with high degrees of functional connectivity that are located medially/centrally in the larval zebrafish forebrain.

**Figure 2.**
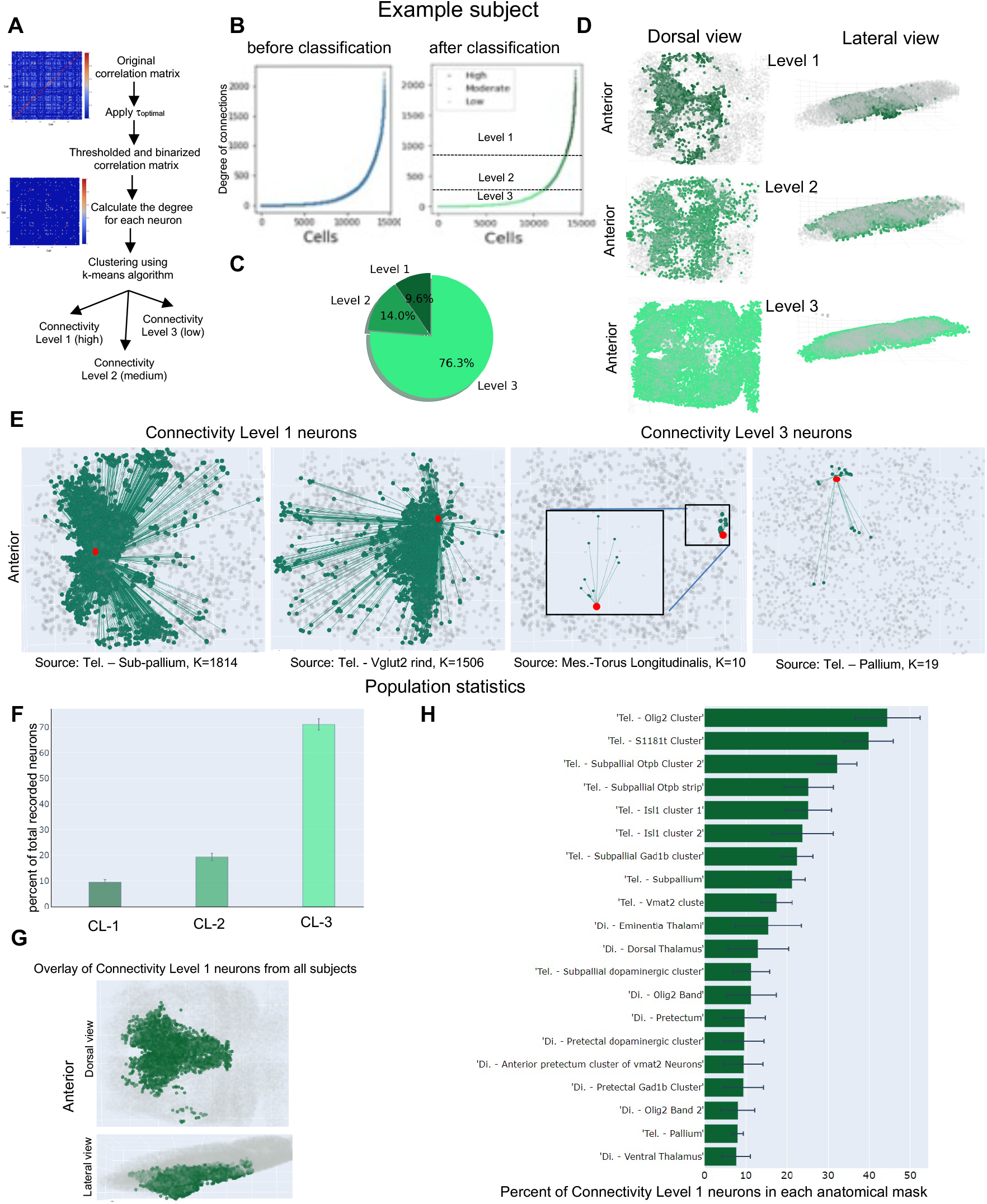
Classification of neurons based on their degree of functional connections. **A,** Overview of the classification of individual ROIs (neurons) based on their level of functional connectivity (degree). The Pearson correlation coefficient was used to calculate the correlation matrix, which was then thresholded using the optimal threshold value. The k-means clustering algorithm was used to cluster ROIs based on their degree. **B,** Sorted ROIs (left) vs. clustered ROIs (right) based on their functional connectivity level for an example subject. **C,** Pie chart showing the percentage of ROIs in three connectivity level categories for an example subject. The ROIs with the highest level of functional connectivity is the smallest group (around than 8%). **D,** Dorsal and lateral views of the three connectivity categories of ROIs’ distributions in the example subject’s forebrain. **E,** The connectivity of ROIs with the connectivity level 1 and 3 in the example subject brain. **F,** Percent of total recorded neurons in each functional connectivity level across 9 subjects. **G,** overlay view of highly functional connected ROIs (Level 1) in all 9 subjects shows that they are located in the medial part of the forebrain. **H,**Anatomical distribution of connectivity Level 1 neurons sorted based on the percentage of total recorded neurons in each anatomical mask.

### Complementary domains of high neuronal activity and high functional connectivity exists in the larval zebrafish forebrain

It was intriguing to note that highly active neurons occupied regions that are complementary to those occupied by highly connected neurons in the larval zebrafish forebrain (Fig. 3A). Plotting the activity and functional connectivity values for all recorded neurons in one example subject showed that highly active neurons did not overlap with highly connective neurons (Fig. 3B). To visualize the distribution and assess statistical significance on a population scale, we used a bootstrapping method (*50*) to construct a graph showing the percentage of neurons with different levels of activity and connectivity at 2.5% and 95% confidence intervals (i.e., AL-1&CL-1, AL-1&CL-2, AL-1&CL-3, AL-2&CL-1, AL-2&CL-2, AL-2&CL-3, AL-3&CL-1, AL-3&CL-2, AL-3&CL-3). Specifically, 9 subjects were randomly sampled with replacement and this was repeated at least 25 times. A non-overlap set of the AL-1 and CL-1 neurons was detected (Fig. 3C), suggesting a mutually exclusive relationship between highly active and highly connected neurons in the larval zebrafish forebrain.

**Figure 3.**
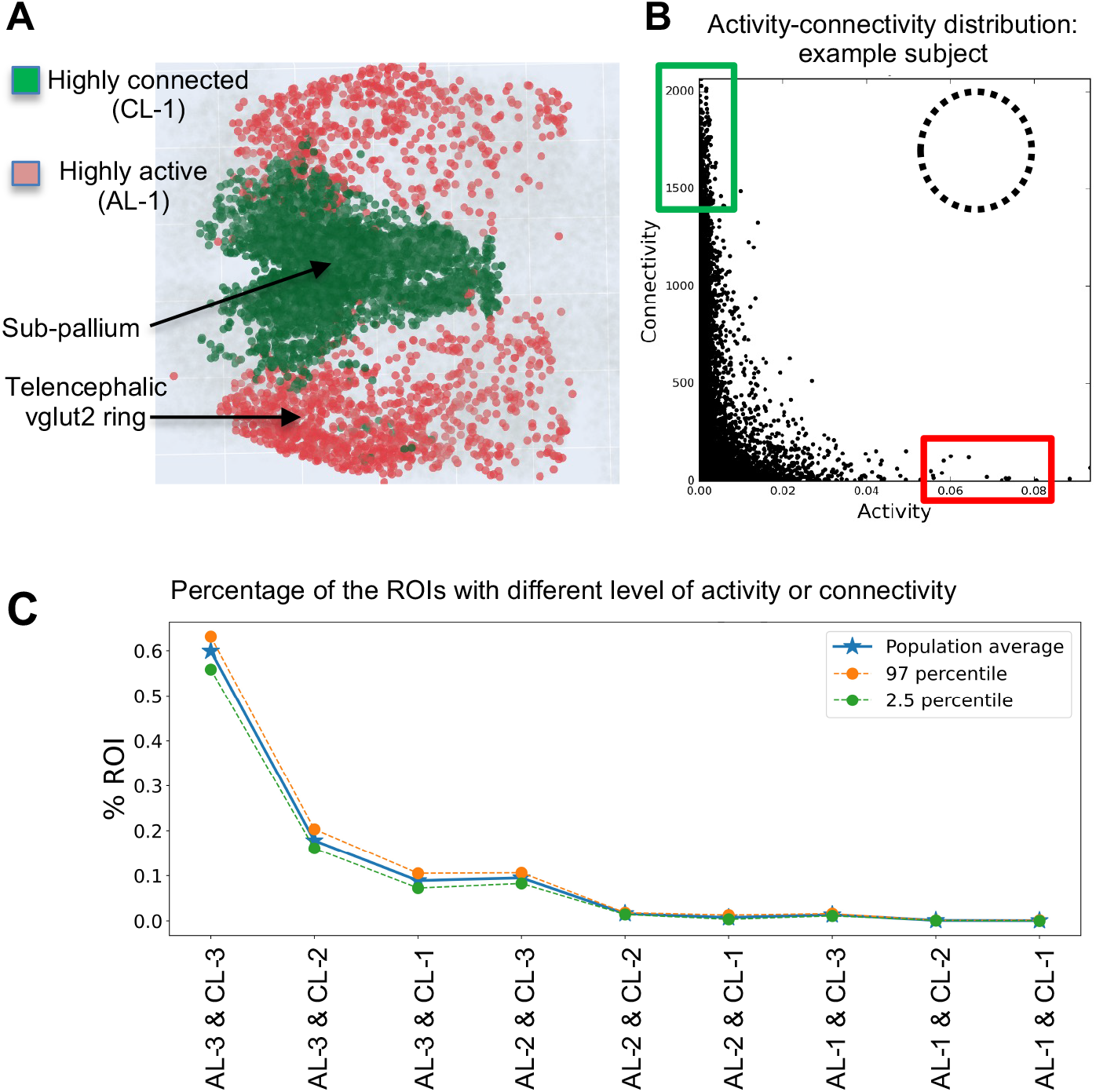
Highly active and highly connected neuronal populations occupy complementary domains in the larval zebrafish forebrain. **A,** Overlay of highly active (red) and highly functional connected ROIs (individual neurons) in the larval zebrafish forebrain across 9 subjects. The highly active cells are in the lateral area whereas the cells with a high level of functional connectivity are located in the medial area of. **B**, connectivity levels (Y-axis) of all neurons sorted based on their activity (X-axis) in an example subject. Red and blue boxes denote neurons of high activity and high functional connectivity respectively. The dotted circle denotes where highly active and highly connected neurons are expected. **C,** the population distribution curve of all neurons with different levels of activity and functional connectivity. Note that neurons that have high activity and high connectivity are non-existent.

### Regions of high neuronal activity versus high functional connectivity are largely non-overlapping in the resting state human brain

To determine whether such relationship between activity and connectivity is an evolutionarily conserved phenomenon, we analyzed the resting state human brain functional magnetic resonance imaging (fMRI) data from Centre for Biomedical Research Excellence (COBRE) dataset (*51*) (http://fcon_1000.projects.nitrc.org/indi/retro/cobre.html)(Fig. 4A). The COBRE data set includes the resting state fMRI data from 74 healthy individuals that were used in this study. The fMRI dataset for each subject includes blood-oxygenation level dependent (BOLD) volumes of 5 minutes. In contrast to the larval zebrafish data in which each ROI is a single neuron, the human fMRI data described activity and connectivity on the scale of brain regions (i.e., each ROI is a brain region). Brodmann areas defined by the Talairach Daemon (TD) system (*52*) were assigned to all subjects (a total of 105 ROIs). Data pre-processing workflow was shown in Fig. S7.

**Figure 4.**
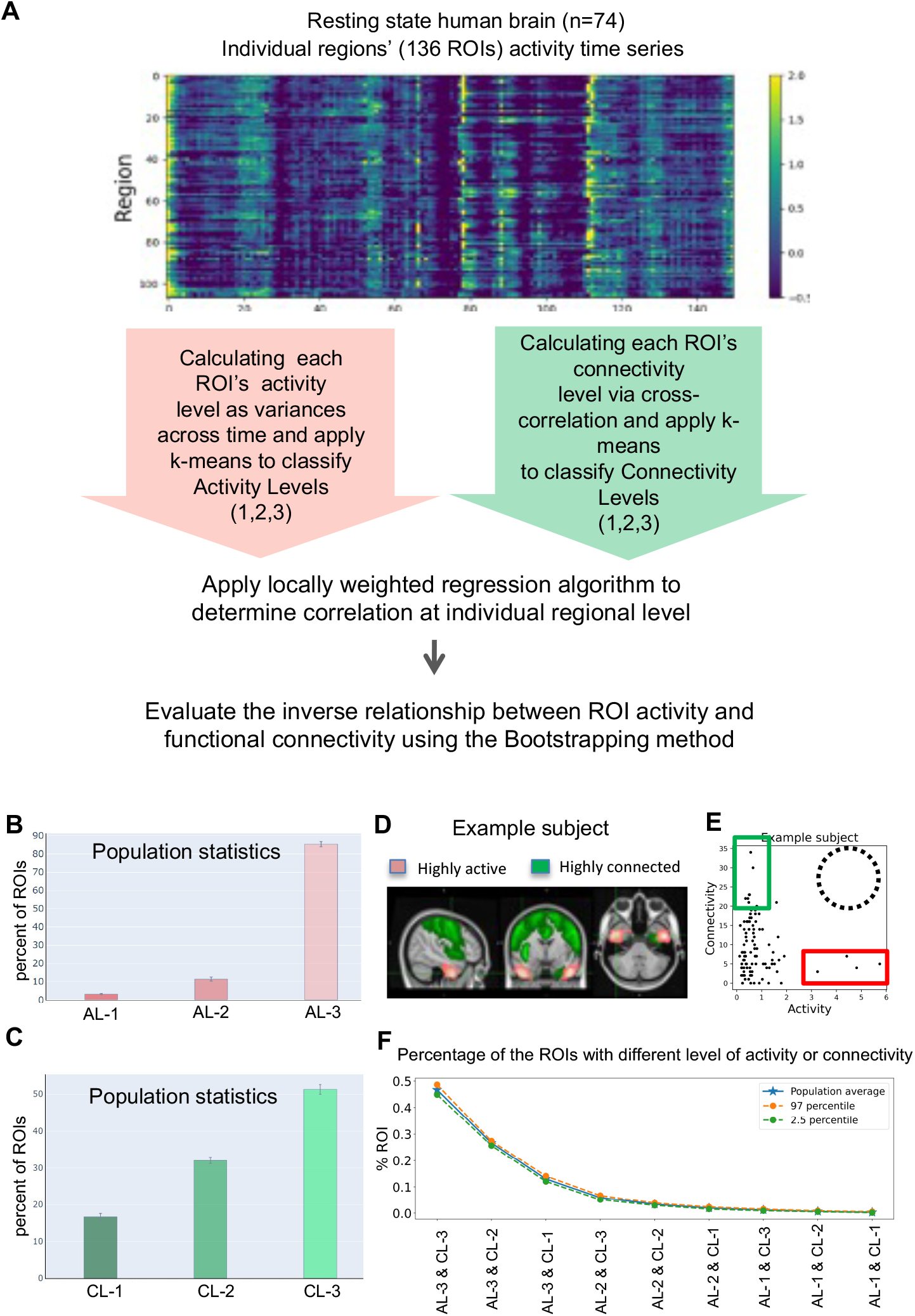
Largely non-overlapping distribution of highly active and highly connected regions in the resting state human brain. **A,** Overview of the classification of individual ROIs (brain regions, n=105) based on their level of activity and functional connectivity (degree). Variations of the brain region activity across time was used as a measure of activity and the Pearson correlation of brain regions’ activity was used a measure of functional connectivity (degree). The k-means clustering algorithm was employed to cluster the brain regions into three levels based on each measure. **B,** Percent of total ROIs in each activity level category. **C,** Percent of total ROIs in each functional connectivity level category. **D,** Highly active (red) and highly connected regions (green) across all subjects (n=74). **E,** connectivity levels (Y-axis) of all brain regions sorted based on their activity (X-axis) in an example subject. Red and blue boxes denote regions of high activity and high functional connectivity respectively. The dotted circle denotes where highly active and highly connected brain regions are expected to locate. **F,** the population distribution curve of brain regions with different levels of activity and functional connectivity. Note that brain regions that have high activity and high connectivity are non-existent.

Similar to our calcium trace data analysis, the variance of BOLD signals across time was calculated as the activity for each ROI, followed by applying the k-means algorithm to classify ROIs into three levels of activity (Fig. 4B, and Fig. S8A-B). Our analysis showed that the anterior division of inferior temporal gyrus displayed high activity that was observed in more than 45% of the subjects (Fig. S8C-D), whereas the hippocampus, pallidum, and putamen were less active in the resting state (Fig. S8E, and Table S1).

We next used the Pearson correlation measure to derive a metric of functional connectivity between ROIs followed by k-means classification into three levels of connections (Fig. 4C, and Fig. S9A-B). Similar to the analysis of functional connectivity in the zebrafish brain, we applied the thresholding value that provided the best fit to the power-law curve as the optimal threshold value for the connectivity matrix. Our analysis showed that the precentral gyrus, right postcentral gyrus and anterior division of Cingulate Gyrus were among the regions with the high level of connectivity in the human brain, whereas the Pallidum right frontal medial cortex, subcallosal cortex, and amygdala were much less connected in almost all subjects in the human resting state dataset (Fig. S9C-E, Table S1).

Similar to the observations made in the larval zebrafish forebrain, highly active regions and highly functionally connected regions in the resting state human brain appear complementary and non-overlapping (Fig. 4D). Plotting the activity and functional connectivity values for all recorded brain regions in one example subject (Fig. 4E) and the percentage of ROIs in each activity and connectivity category (Fig. 4F) further reinforced this notion. Together, regions with high functional connectivity appear mutually exclusive with those of high activity in the resting state human brain.

## Discussion

One major goal of neuroscience is to understand fundamental organizational principles of the brain. While functional imaging and analysis of brain networks in larval zebrafish is an emerging field, numerous studies of resting state human brain networks have examined brain activity or connectivity patterns, suggesting the prevalence of activity-based or connectivity-based organizations (*35, 53*).

Despite these advances, the relationship between activity and functional connectivity in the brain has not been previously examined. This is an interesting and important question both for understanding the brain architectural principles and for designing artificial neuronal networks. In this study, we have examined the relationship between activity and functional connectivity in both the larval zebrafish forebrain where each ROI is an individual neuron, and in the resting state human brain where each ROI is a brain region composed of millions of neurons. In larval zebrafish, activity is measured through quantifying variances of calcium signals over time: more frequent events of fluorescent changes indicate higher activity. In the human brain, activity is measured through the BOLD signal, i.e., alterations in deoxyhemoglobin driven by localized changes in brain blood flow and blood oxygenation, which are coupled to underlying neuronal activity. More frequent events of BOLD signals indicate higher activity. Functional connectivity in both the larval zebrafish and human brains is measured using Pearson correlation; the resulting correlation matrices are further denoised with optimal threshold values, which are determined using the concept of “small-world” network with power law distribution. Through these analyses, we have uncovered a mutually exclusive relationship between high activity and high functional connectivity at individual ROI levels across all zebrafish and human subjects. This is remarkable, given the 450 million years of evolutionary distance and the drastic brain size differences (100K vs. 100 billion neurons) between the two species.

We propose two possible models to explain why such exclusive relationship between high activity and high functional connectivity is at work in both zebrafish and human brains. The first model pertains to a physical constraint. Given that structural and functional connectivity show considerable correlation (*16*), it is possible that neurons with high levels of connections are physically incapable of achieving high activity. The second model is based on a metabolic constraint. ROIs with high activity are at a high metabolic cost (*54*), thereby accumulating more oxidative damage and prone to degeneration. To best preserve network integrity, it would therefore be desirable to delegate the tasks that require high activity to the ROIs with low degree of functional connections, while maintaining ROIs with high connections at low activity. Future experiments are necessary to test these models. With the accessibility of the zebrafish brain to molecular cellular and systems level dissections, such validations are feasible in zebrafish.

## Acknowledgments

We thank Misha Ahrens for providing the transgenic line *Tg[HuC-H2B-GCAMP6s]*, Ryan McGorty for constructing the iSPIM microscope, Saul Kato and other Guo lab members for helpful discussions, Michael Munchua and Vivian Yuan for excellent animal care. This project was supported by NIH R01 GM132500, R21 DA044007, and R21 MH113961 to S.G.

## Author contributions

M.Z., D.X. and S.G. designed the experiments and interpreted the results, D.X. performed the calcium imaging experiments and calcium image data processing, M.Z. performed data analysis with the assistance from F.J., A.A., B.H., S.N., A.R. M.Z., D.X, and S.G. wrote the manuscript, with the input from all authors.

## Materials and Correspondence

Correspondence and material requests should be addressed to M.Z or S.G.

## Supplementary Info

### METHODS

#### Zebrafish strain maintenance and larval sample preparation

The transgenic line *Tg[HuC-H2B-GCaMP6s]* with *nacre* or *casper* background was used for breeding. Embryos were kept in blue egg water (2.4 g of CaSO4, 4g of Instant Ocean Salts, 600μl of 1% Methylene blue in 20 liters of milliQ water) and incubated at 28°C. On 6 days post fertilization (dpf), healthy larvae with high GCaMP6s expression were selected for imaging. Fish samples are held in custom-designed polydimethylsiloxane (PDMS) sample holders with each holder carrying up to 5 larvae. Each larva was half-embedded in a slot on the sample holder, paralyzed with 1mg/mL mivacurium chloride and covered with 2% agarose gel/E3 medium solution. After loading the whole group of larvae to be imaged, the sample holders were immersed in E3 medium for 1 hour to wash off traces of mivacurium chloride. All animal experiments were approved by the Institutional Animal Care and Use Committee (IACUC) at the University of California, San Francisco, USA.

#### *In vivo* calcium imaging using iSPIM

An inverted SPIM (iSPIM) that is similar to a previously reported design (*39*) was used for imaging. The microscope framework was adapted from a di-SPIM (Applied Scientific Imaging, Inc.) (*55*). The excitation laser (Coherent OBIS LS 488 nm) was fiber-coupled into the microscope, collimated, then focused by a 0.3 NA water-dipping objective (Nikon). A virtual light-sheet was created by deflecting the scanning mirror, illuminating a layer of specimen with ~7 μm thickness. Fluorescence from the illuminated layer was collected in an orthogonal direction by a 20x 1.0 NA water-dipping objective (Olympus XLUMPLFLN-W) and the final image is captured by a scientific CMOS (sCMOS) camera (Hamamatsu Orca Flash 4.0, C11440). Due to the strong scattering of the blue-green light in deep tissues, we confined the illumination within dorsal forebrain of the larvae. The volume of interest was approximately 300 μm × 250 μm x 100μm in x,y,z directions, respectively. This volume covered the entire dorsal telencephalon and habenula regions. To resolve the dynamics of GCaMP6s, image stacks were acquired at 2Hz, which allowed us to resolve frequency components up to 1Hz based on Nyquist sampling theorem. Each 100μm stack consisted of 26 slices with 4 μm between two adjacent slices. The resting state of the selected volume in each larva was imaged for 15 minutes; in addition to this time lapse recording, a Z-stack with 1 μm step (101 slices) was acquired across the same volume as a reference. The raw data were submitted to the data preprocessing pipeline for cleaning and feature extraction.

#### Pre-processing of calcium imaging data

##### Drift correction and ROI (neuron) extraction

Raw images were organized as hyper stacks in the order of x-y-z-t. Each hyper stack was split into 26T-stacks at different z positions and drift corrected with the StackReg plugin(*56*). For each drift-corrected T-stack, neuronal nuclei were segmented using the Laplacian of gaussian blob detection algorithm blob_log in the Python library scikit image (*57*). Since neuronal nuclei (~4μm in diameter) can be imaged in two or more adjacent planes, a redundancy detection algorithm was developed to find lateral duplications in cell extraction: if a nucleus is detected at the same (x; y) position in the kth and the (k + 1)th planes, this detection would be considered as redundant and the Z-position was considered as an average between zk and zk+1. In each extracted neuronal nuclei, its raw fluorescence signals were calculated through the entire T-stack, and the relative signal intensity of calcium transients, ΔF/F, was calculated using the method as previously described (*45*). Two additional cleaning steps were applied to remove potential artifacts that were falsely recognized as neuronal nuclei by the blob detection algorithm: first, since GCaMP6s has background signals in the absence of action potentials, blobs of real neurons should have a high fluorescence baseline, and blobs with very low baseline (comparable to dark areas in the image) are excluded; second, since in reality the value of ΔF/F should fall within a reasonable range, blobs with extraordinarily high ΔF/F values (exceeding a threshold during recording) were excluded.

Since activity levels vary among neurons, over the 15-min imaging session, some neurons may exhibit high calcium signal peaks while others remain “silent”. Since the latter are not likely to contribute to downstream analyses, it would be beneficial to exclude them from the very beginning. This requires us to: 1). find a reliable method to identify peaks from background in a noisy timeseries; 2). find a measure for the activity level of each neuron, i.e., whether and how much does it activate during the imaging session; 3). Set a well-defined criterion to accept or reject a neuron based on its two characteristics above. The method we used to identify peaks and baselines from each ΔF/F time trace was based on the Bayesian inference of two-dimensional distribution of adjacent ΔF/F values (*31*). For each neuron i, its baseline of ΔF/F, μ_i_, was calculated by averaging all the (ΔF/F)_I_ time points that are identified as background,

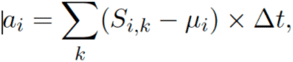

where S_i,k_ refers to the kth time point of (ΔF/F)_i_.

##### Image registration

Since individuals differ in morphology, placement, and brightness, it is difficult to directly compare their images even though the latter are acquired under the same condition. In order to compare imaging results from different individuals, the images should be anatomically mapped to a common template image, i.e., a “reference”. A whole-brain template, Z-brain, has been provided by Randlett et al (*44*) as a standard reference brain atlas for anatomical and functional studies of larval zebrafish brain. Although the fish we experimented on were at the same stage as that in the Z-brain template, the iSPIM stacks acquired in their dorsal forebrain could not be directly registered to the reference due to: 1) a significant mismatch between two imaging directions and 2) a significant difference between the volumes of interests. Since the dorsal forebrain region that we imaged only accounts for a sub-volume of the entire brain, a direct registration of the former to the latter is prone to error. To solve this problem, we created an intermediate reference brain from the Z-brain template by resampling the dorsal-forebrain region in the direction and pixel sizes that are comparable to those of the iSPIM stacks. For each fish, its densely sampled Z-stack was registered to the intermediate reference brain using computational morphometry toolkit (CMTK) (http://nitrc.org/projects/cmtk). After registering the iSPIM stacks into the intermediate template, the coordinates of the extracted neurons in original T-stacks need to be reformatted accordingly into the registered frame. This step was carried out using the registration output and the function stream x form in CMTK. By comparing the reformatted coordinates with the 294 anatomical masks in the Z-brain template, we were able to identify the anatomical label of each neuron and the brain regions covered by our imaging volume.

#### Preprocessing of the resting state human brain fMRI data

We used healthy control subjects (n=74) from the COBRE^16^ resting-state fMRI datasets available at http://fcon_1000.projects.nitrc.org/indi/retro/cobre.html. The fMRI dataset for each subject includes blood-oxygenation level-dependent (BOLD) volumes of 5 minutes (TR = 2 s, TE = 29 ms, FA = 75°, 32 slices, voxel size = 3×3×4 mm^3^, matrix size = 64×64, FOV = 255 ×255 mm^2^). The pre-processing steps included realignment, co-registering, and normalization. We used established preprocessing and analysis pipelines (*58*) and CONN software package (https://www.nitrc.org/projects/conn).

##### Re-alignment

In brief, the first-level covariate containing the 6 rigid-body parameters was created based on the MRI data to estimate the subject motion. For each subject, this variable was used to perform regression on the fMRI data to correct for motion-related effects.

To reduce the physiological noise source, a Component-Based Noise Correction Method (CompCor) was used (*59*).

##### Co-registering

The functional volumes are co-registered with the ROIs and structural volumes. All the Brodmann areas (ROIs) defined through the Talairach Daemon (TD) system (*52*) were assigned to all subjects using segmentation of structural image; grey matter, white matter and cerebrospinal fluid (CSF) masks were generated. Anatomical volumes were co-registered to the functional and ROI volumes for each subject and the volumes were transformed to the MNI-space.

##### Calculation and normalization of fMRI measures

Following re-alignment and co-registering, the Principal Component Analysis (PCA) algorithm was used to extract BOLD signal components for each ROI. The fMRI measures were calculated using MATLAB-based software packages, SPM12 (http://www.fil.ion.ucl.ac.uk/spm/). All of the computed measures are normalized to an N (0,1) Gaussian distribution for each subject.

#### Data analysis

##### Analysis and classification of ROI activity levels

For a given ROI (i.e., individual neurons in larval zebrafish or brain regions in the case of human fMRI data) *i*, if *s*_*i*_ is denoted as its variance of ΔF/F (zebrafish) or BOLD (human) signals over time, the activity value *ac*_*i*_, which is the mean of squared deviations from the mean of *s*_*i*_, was calculated as follows:

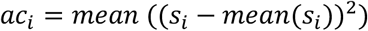

After obtaining each ROI activity level in the given time series, we used the K-means algorithm (*47*) to classify them into Activity Levels 1–3. The k-means clustering algorithm minimizes the within-cluster squared Euclidean distances. Here a one-dimensional activity population was partitioned into 3 sets (levels). The within-class cells in each level have similar activity.

##### Analysis and classification of ROI functional connectivity levels

We used Pearson correlation to measure functional connectivity between ROIs. Since Pearson correlation assigns a value to all ROI pairs, it is necessary to apply thresholding to eliminate potentially spurious connections. There is no standard method to calculate the optimal threshold value *τ*_*optimal*_ and different values of *τ* are used to create the adjacency matrices. Arbitrarily chosen thresholding values are often applied to raw matrixes (*6*). As different cutoff values can directly influence network properties and bias analysis results, we developed algorithms to calculate the optimal thresholding values and generate the connectivity matrix. This matrix was then used to calculate the functional connections (i.e., degrees) of each ROI, followed by K-means classification into three connectivity levels. The steps of calculating functional connections for each ROI are:

1. Let *ρ* = *ρ*_*ij*_ be the correlation matrix, where *ρ*_*ij*_ is the Pearson correlation of ROIs *i* and *j* and can be calculated as follows:

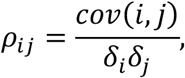
2. Setting the threshold values:

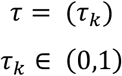
3. Thresholding the connectivity matrix using *τ*_*k*_ ∈ *τ* or each *ρ*_*ij*_

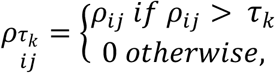

where 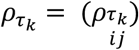 is the thresholded correlation matrix.
4. Binarizing 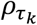

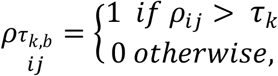
5. Calculating the degrees for each *ROI*_*i*_, *degree*_*i*_ = ∑_*j*_*l*_*i,j*_, where *l*_*i,j*_ is the link between *roi*_*i*_ and *roi*_*j*_.
6. Calculating the degree distribution of the network; The fraction of ROIs with the degree k is defined as follows:

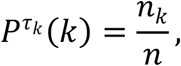

where n_k_, is the number of the ROIs that have degree k.
7. Calculating the fitting value of the degree distribution with the power-law distribution. *r*^2^ (coefficient of determination) was used to evaluate the closeness of data at each threshold value to the power-law curve.
8. Derive the optimal threshold value: At the optimal threshold value, the *r*^2^ is highest, which indicates the best fit of the data to the power-law curve.

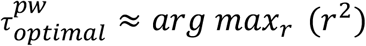

The detailed steps of calculating the optimal threshold value of the connectivity matrix were provided in the Fig. S4. The average 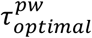 value of all zebrafish subjects in our data ranged from 0.4 to 0.6. We applied this algorithm to the human fMRI data and obtained 0.7 for 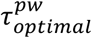. Using these values, we binarized our data: correlations with a value less than 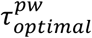 were set to zero, whereas those with a value greater than 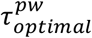 were set to one.

To further test whether the observed power-law structure of the functional brain is relevant, we shuffled the data (Fig. S5) to generate random networks with the same numbers of nodes and edges as the original networks and applied similar thresholding and analysis of degree distributions. The random network did not follow a power law structure at any thresholding values tested. Together, these analyses enable us to establish optimal thresholding values that uncover biologically relevant networks. Using such matrixes, we were able to detect known connections between the olfactory epithelia and the olfactory bulb (Fig. S6).

After obtaining each ROI’s numbers of functional connections in the given time series, we used the K-means algorithm as described above to classify them into Connectivity Levels 1–3.

##### Analysis of the relationship between activity and functional connectivity

We plotted the activity and connectivity for each ROI in each individual. The input data are: zebrafish calcium imaging data are composed of >12k individual neuronal activity and connectivity per subject (n=9 subjects); human resting-state fMRI data are composed of 136 brain regions’ activity and connectivity per subject (n=74 subjects).

To analyze the population frequency of each class of ROIs (i.e., AL-1&CL-1, AL-1&CL-2, AL-1&CL-3, AL-2&CL-1, AL-2&CL-2, AL-2&CL-3, AL-3&CL-1, AL-3&CL-2, AL-3&CL-3) and its statistical significance, we used the following bootstrapping method:

1. Select the number of the bootstrapping iteration (here, 25 iterations were used).
2. Repeat the steps “3” to “7” 25 times for the input data.
3. Select a sample set with replacement from the set of all subjects.
4. Calculate activity (variances) and connectivity (degrees) data for each ROI from all sampled subjects.
5. Calculate activity (variances) and connectivity (degrees) data for each ROI from all sampled subjects.
6. Calculate the average ROI population size for each activity and connectivity level.
7. Calculate the mean, lower (2.5 percentile), and upper (97.5 percentile) point-wise confidence bands for the populations that are calculated in the step “6”.

#### Data visualization

Different python libraries were used to visualize the results. The plotly libraries (https://plotly.com/python/) were used to visualize the cells’ anatomical and spatial distributions in the calcium imaging data. We used the FSL (https://fsl.fmrib.ox.ac.uk/fsl) to visualize the brain regions in the fMRI data.

#### Statistical analysis

Sample sizes and statistics are reported in the figure legends and text for each measurement. To determine the relationship between activity and functional connectivity at individual ROI levels (Fig. 3c and 4e), we used bootstrap tests (with 7 iterations) to test whether the negative correlation between neuronal activity and functional connectivity is consistent across subjects. For each subject, the activity (variances) and connectivity (degrees) for each ROI were calculated. Individual neurons in the larval zebrafish calcium imaging data and individual brain regions in the human fMRI data were considered as ROIs. The ROIs were sorted based on their activity for each subject (X-axis). The connectivity of sorted ROIs for each subject (Y-axis) was then graphed. The Locally Weighted Regression algorithm (*60*) was applied for each subject’s data to approximate the polynomial curve. Finally, the bootstrapping method (*50*) was used to evaluate the inverse relationship between neuronal activity and functional connectivity. Here random sampling with replacement was applied for selecting the curves fitting of individual subjects to evaluate the model.

## Supplementary Info

### METHODS

#### *In vivo* calcium imaging using iSPIM

An inverted SPIM (iSPIM) that is similar to a previously reported design (*1*) was used for imaging. The microscope framework was adapted from a di-SPIM (Applied Scientific Imaging, Inc.) (*2*). The excitation laser (Coherent OBIS LS 488 nm) was fiber-coupled into the microscope, collimated, then focused by a 0.3 NA water-dipping objective (Nikon). A virtual light-sheet was created by deflecting the scanning mirror, illuminating a layer of specimen with ~7 μm thickness. Fluorescence from the illuminated layer was collected in an orthogonal direction by a 20x 1.0 NA water-dipping objective (Olympus XLUMPLFLN-W) and the final image is captured by a scientific CMOS (sCMOS) camera (Hamamatsu Orca Flash 4.0, C11440). Due to the strong scattering of the blue-green light in deep tissues, we confined the illumination within dorsal forebrain of the larvae. The volume of interest was approximately 300 μm x 250 μm x 100μm in x,y,z directions, respectively. This volume covered the entire dorsal telencephalon and habenula regions. To resolve the dynamics of GCaMP6s, image stacks were acquired at 2Hz, which allowed us to resolve frequency components up to 1Hz based on Nyquist sampling theorem. Each 100μm stack consisted of 26 slices with 4 μm between two adjacent slices. The resting state of the selected volume in each larva was imaged for 15 minutes; in addition to this time lapse recording, a Z-stack with 1 μm step (101 slices) was acquired across the same volume as a reference. The raw data were submitted to the data preprocessing pipeline for cleaning and feature extraction.

#### Pre-processing of calcium imaging data

##### Drift correction and ROI (neuron) extraction

Raw images were organized as hyper stacks in the order of x-y-z-t. Each hyper stack was split into 26T-stacks at different z positions and drift corrected with the StackReg plugin(*3*). For each drift-corrected T-stack, neuronal nuclei were segmented using the Laplacian of gaussian blob detection algorithm blob_log in the Python library scikit image (*4*). Since neuronal nuclei (~4μm in diameter) can be imaged in two or more adjacent planes, a redundancy detection algorithm was developed to find lateral duplications in cell extraction: if a nucleus is detected at the same (x; y) position in the kth and the (k + 1)th planes, this detection would be considered as redundant and the Z-position was considered as an average between zk and zk+1. In each extracted neuronal nuclei, its raw fluorescence signals were calculated through the entire T-stack, and the relative signal intensity of calcium transients, ΔF/F, was calculated using the method as previously described (*5*). Two additional cleaning steps were applied to remove potential artifacts that were falsely recognized as neuronal nuclei by the blob detection algorithm: first, since GCaMP6s has background signals in the absence of action potentials, blobs of real neurons should have a high fluorescence baseline, and blobs with very low baseline (comparable to dark areas in the image) are excluded; second, since in reality the value of ΔF/F should fall within a reasonable range, blobs with extraordinarily high ΔF/F values (exceeding a threshold during recording) were excluded.

Since activity levels vary among neurons, over the 15-min imaging session, some neurons may exhibit high calcium signal peaks while others remain “silent”. Since the latter are not likely to contribute to downstream analyses, it would be beneficial to exclude them from the very beginning. This requires us to: 1). find a reliable method to identify peaks from background in a noisy timeseries; 2). find a measure for the activity level of each neuron, i.e., whether and how much does it activate during the imaging session; 3). Set a well-defined criterion to accept or reject a neuron based on its two characteristics above. The method we used to identify peaks and baselines from each ΔF/F time trace was based on the Bayesian inference of two-dimensional distribution of adjacent ΔF/F values (*6*). For each neuron i, its baseline of ΔF/F, μ_i_, was calculated by averaging all the (ΔF/F)_I_ time points that are identified as background,

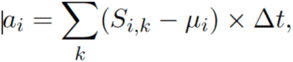

where S_i,k_ refers to the kth time point of (ΔF/F)_i_.

##### Image registration

Since individuals differ in morphology, placement, and brightness, it is difficult to directly compare their images even though the latter are acquired under the same condition. In order to compare imaging results from different individuals, the images should be anatomically mapped to a common template image, i.e., a "reference". A whole-brain template, Z-brain, has been provided by Randlett et al (*7*) as a standard reference brain atlas for anatomical and functional studies of larval zebrafish brain. Although the fish we experimented on were at the same stage as that in the Z-brain template, the iSPIM stacks acquired in their dorsal forebrain could not be directly registered to the reference due to: 1) a significant mismatch between two imaging directions and 2) a significant difference between the volumes of interests. Since the dorsal forebrain region that we imaged only accounts for a sub-volume of the entire brain, a direct registration of the former to the latter is prone to error. To solve this problem, we created an intermediate reference brain from the Z-brain template by resampling the dorsal-forebrain region in the direction and pixel sizes that are comparable to those of the iSPIM stacks. For each fish, its densely sampled Z-stack was registered to the intermediate reference brain using computational morphometry toolkit (CMTK) (http://nitrc.org/projects/cmtk). After registering the iSPIM stacks into the intermediate template, the coordinates of the extracted neurons in original T-stacks need to be reformatted accordingly into the registered frame. This step was carried out using the registration output and the function stream x form in CMTK. By comparing the reformatted coordinates with the 294 anatomical masks in the Z-brain template, we were able to identify the anatomical label of each neuron and the brain regions covered by our imaging volume.

#### Preprocessing of the resting state human brain fMRI data

We used healthy control subjects (n=74) from the COBRE^16^ resting-state fMRI datasets available at http://fcon_1000.projects.nitrc.org/indi/retro/cobre.html. The fMRI dataset for each subject includes blood-oxygenation level-dependent (BOLD) volumes of 5 minutes (TR = 2 s, TE = 29 ms, FA = 75°, 32 slices, voxel size = 3×3×4 mm^3^, matrix size = 64×64, FOV = 255 ×255 mm^2^). The pre-processing steps included realignment, co-registering, and normalization. We used established preprocessing and analysis pipelines (*8*) and CONN software package (https://www.nitrc.org/projects/conn).

##### Re-alignment

To reduce the physiological noise source, a Component-Based Noise Correction Method (CompCor) was used (*9*).

##### Co-registering

The functional volumes are co-registered with the ROIs and structural volumes. All the Brodmann areas (ROIs) defined through the Talairach Daemon (TD) system (*10*) were assigned to all subjects using segmentation of structural image; grey matter, white matter and cerebrospinal fluid (CSF) masks were generated. Anatomical volumes were co-registered to the functional and ROI volumes for each subject and the volumes were transformed to the MNI-space.

#### Data analysis

##### Analysis and classification of ROI activity levels

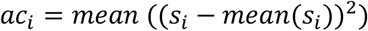

After obtaining each ROI activity level in the given time series, we used the K-means algorithm (*11*) to classify them into Activity Levels 1–3. The k-means clustering algorithm minimizes the within-cluster squared Euclidean distances. Here a one-dimensional activity population was partitioned into 3 sets (levels). The within-class cells in each level have similar activity.

##### Analysis and classification of ROI functional connectivity levels

We used Pearson correlation to measure functional connectivity between ROIs. Since Pearson correlation assigns a value to all ROI pairs, it is necessary to apply thresholding to eliminate potentially spurious connections. There is no standard method to calculate the optimal threshold value *τ*_*optimal*_ and different values of *τ* are used to create the adjacency matrices. Arbitrarily chosen thresholding values are often applied to raw matrixes (*12*). As different cutoff values can directly influence network properties and bias analysis results, we developed algorithms to calculate the optimal thresholding values and generate the connectivity matrix. This matrix was then used to calculate the functional connections (i.e., degrees) of each ROI, followed by K-means classification into three connectivity levels. The steps of calculating functional connections for each ROI are:

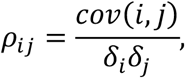
2. Setting the threshold values:

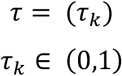
3. Thresholding the connectivity matrix using *τ*_*k*_ ∈ *τ* or each *ρ*_*ij*_

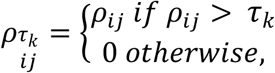

where 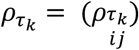 is the thresholded correlation matrix.
4. Binarizing 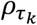

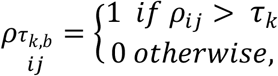
5. Calculating the degrees for each *ROI*_*i*_, *degree*_*i*_ = ∑_*j*_*l*_*i,j*_, where *l*_*i,j*_ is the link between *roi*_*i*_ and *roi*_*j*_.
6. Calculating the degree distribution of the network; The fraction of ROIs with the degree k is defined as follows:

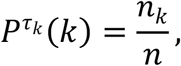

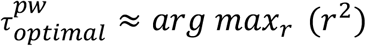

The detailed steps of calculating the optimal threshold value of the connectivity matrix were provided in the Fig. S4. The average 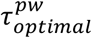 value of all zebrafish subjects in our data ranged from 0.4 to 0.6. We applied this algorithm to the human fMRI data and obtained 0.7 for 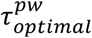. Using these values, we binarized our data: correlations with a value less than 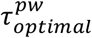 were set to zero, whereas those with a value greater than 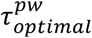 were set to one.

#### Statistical analysis

Sample sizes and statistics are reported in the figure legends and text for each measurement. To determine the relationship between activity and functional connectivity at individual ROI levels (Fig. 3c and 4e), we used bootstrap tests (with 7 iterations) to test whether the negative correlation between neuronal activity and functional connectivity is consistent across subjects. For each subject, the activity (variances) and connectivity (degrees) for each ROI were calculated. Individual neurons in the larval zebrafish calcium imaging data and individual brain regions in the human fMRI data were considered as ROIs. The ROIs were sorted based on their activity for each subject (X-axis). The connectivity of sorted ROIs for each subject (Y-axis) was then graphed. The Locally Weighted Regression algorithm (*13*) was applied for each subject’s data to approximate the polynomial curve. Finally, the bootstrapping method (*14*) was used to evaluate the inverse relationship between neuronal activity and functional connectivity. Here random sampling with replacement was applied for selecting the curves fitting of individual subjects to evaluate the model.

---

**Figure S1.**
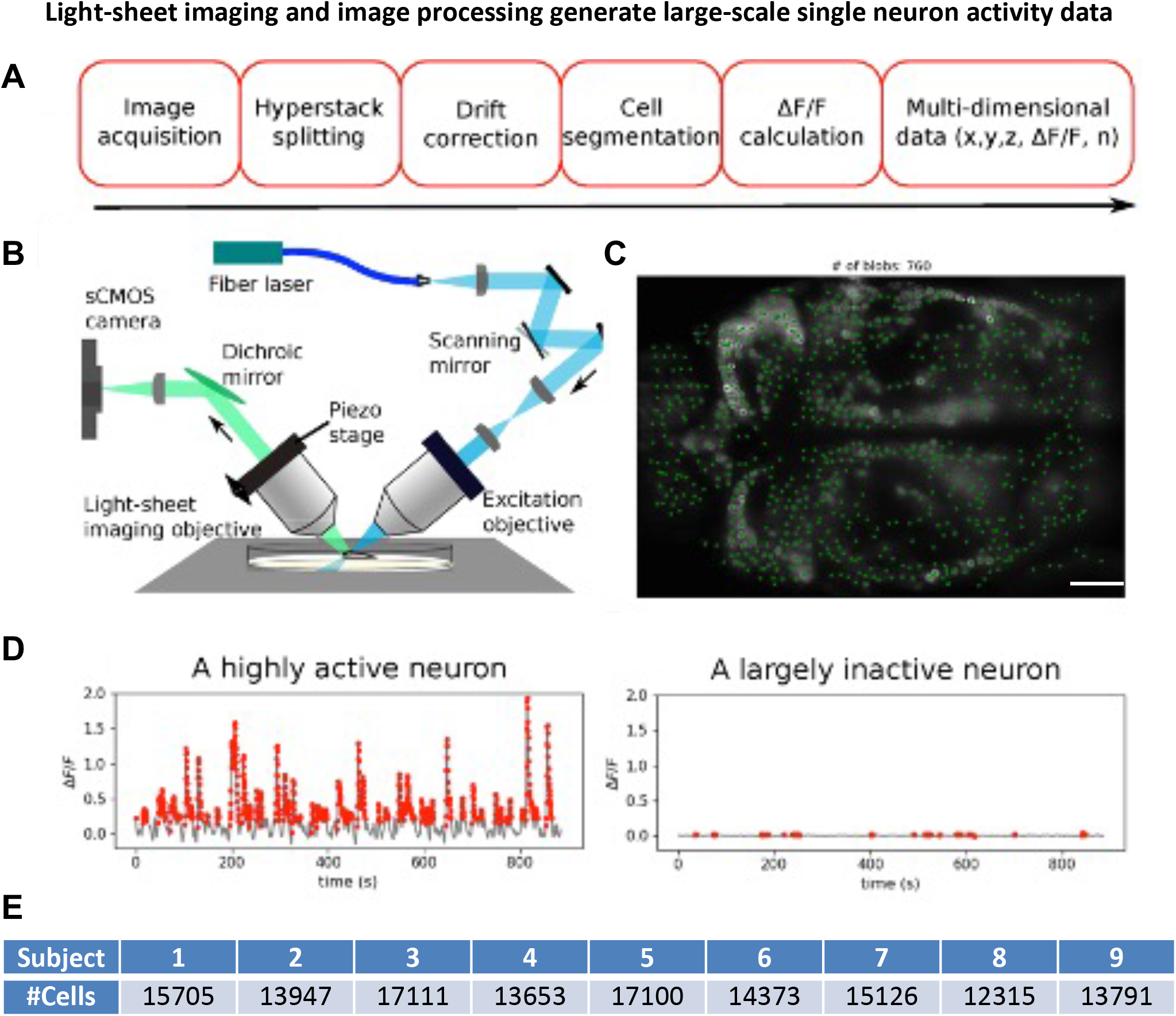
Light-sheet imaging and image processing generate large-scale single neuron activity data. **A**, a schematic showing the overall workflow of image acquisition and pre-processing pipeline resulting in a multi-dimensional dataset. The df/f, 3d coordinate, and anatomical mask was assigned to each detected cell. **B,** a schematic showing the setup of the iSPIM *in vivo* calcium imaging system. **C,** a representative image of 6 dpf larval zebrafish forebrain showing detected cells (blobs) (n=760 cells detected). **D,** An example of a highly active (left) and a largely inactive (right) neurons. **E,** Number of detected cells in each of the 9 subjects used in this study. scale bar, 25 um; dpf, days post fertilization.

**Figure S2.**
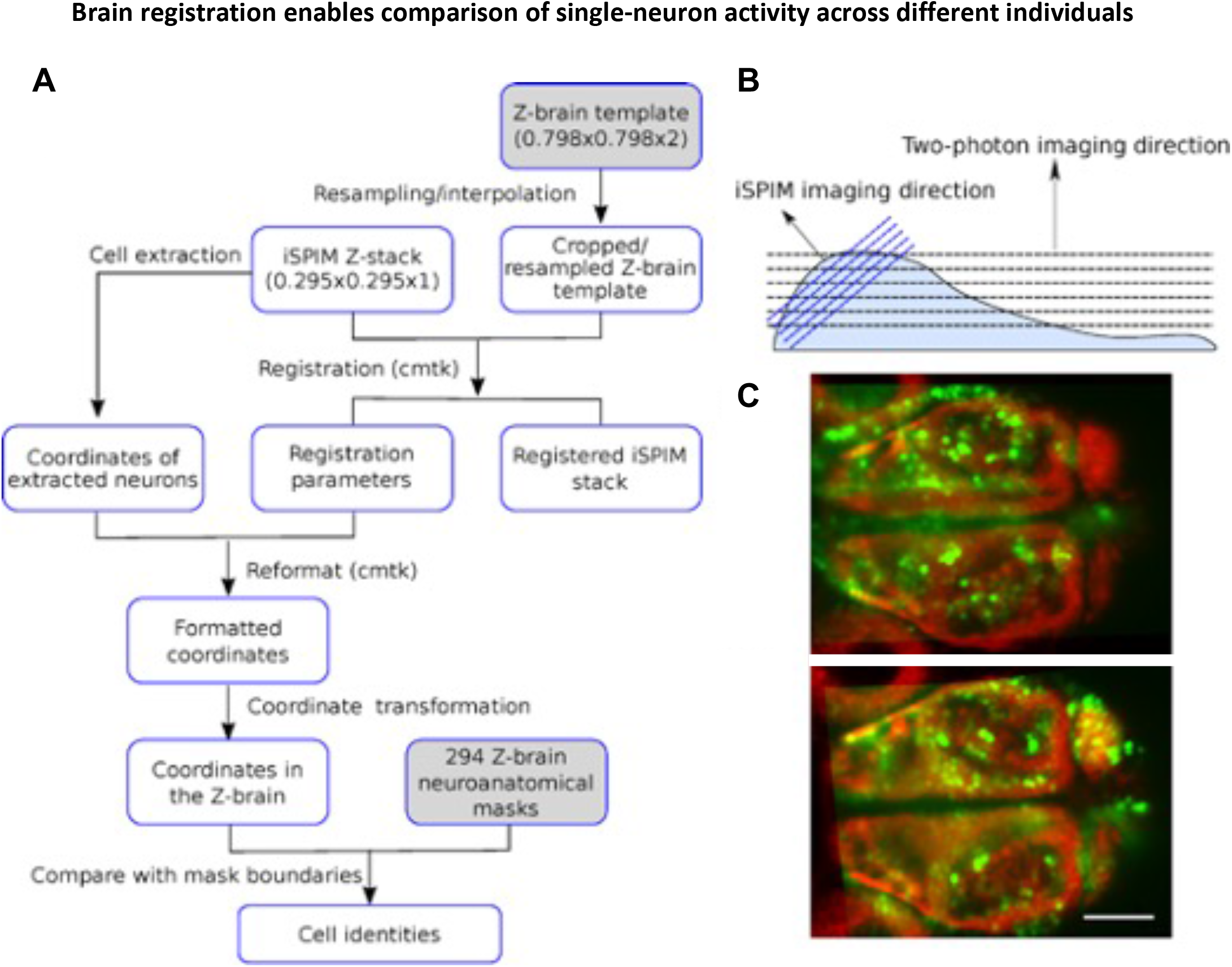
Brain registration enables comparison of single-neuron activity across different individuals. **A,** a schematic showing the workflow of image registration and anatomical labeling. **B,** a schematic showing the imaging direction in the iSPIM (blue) and the 2-photon system (black), respectively. Stacks acquired from the former need to be rotated during image registration to the Z-brain atlas, which is acquired via 2-photon microscopy. **C**, Example image slice of dorsal forebrain, before (top) and after (bottom) registration. Red: 2-photon image of HuC-H2B-RFP labeled forebrain template in the Z-brain atlas, cropped and resampled in the iSPIM imaging direction. Green: HuC-H2B-GCAMP6s labeling. Scale bar, 50 um.

**Figure S3.**
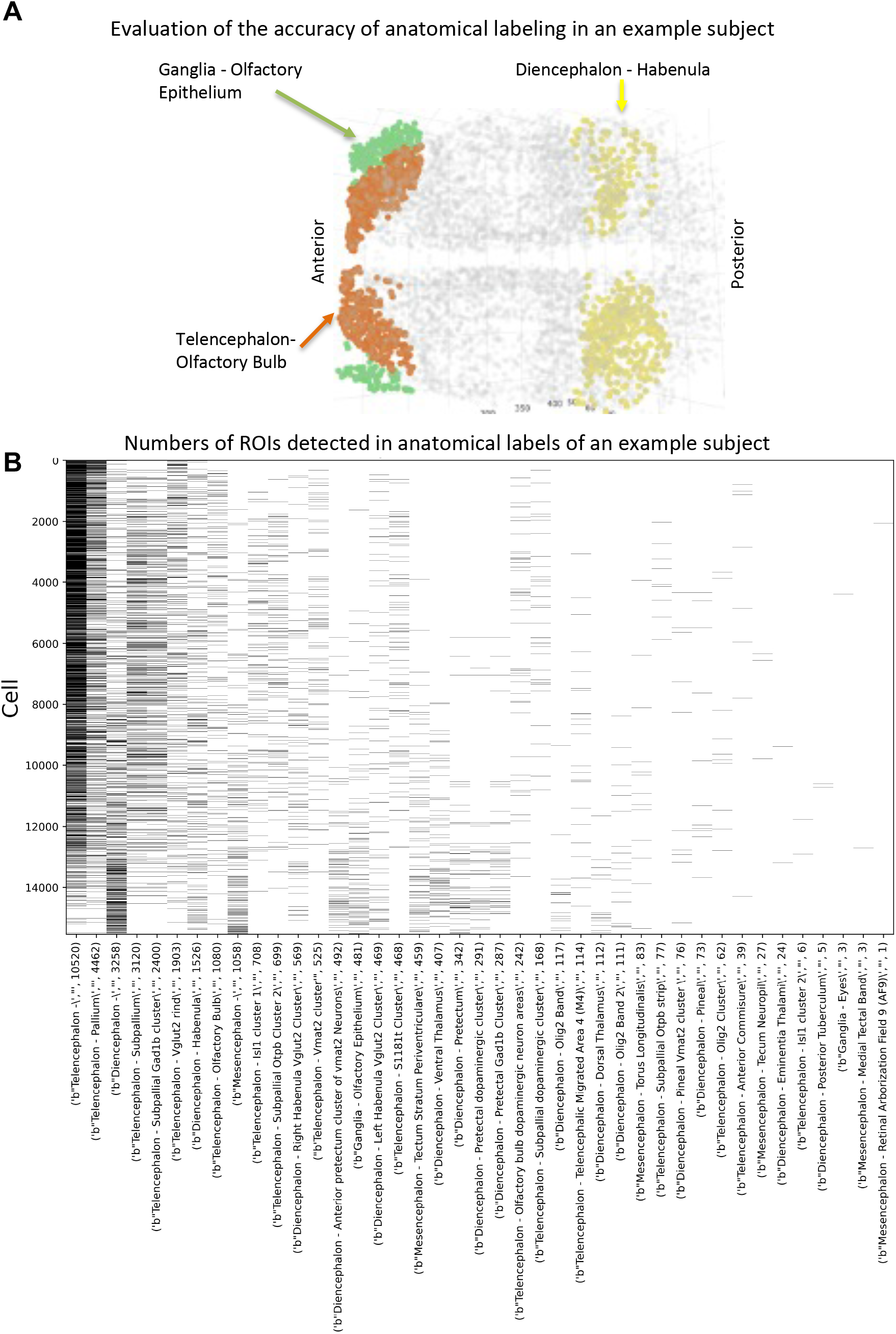
Evaluation of the accuracy and representation of anatomical label assignment. **A,** 2-D visualization of recorded ROIs (individual neurons) in several representative anatomic regions of an example subject. **B,** a binary heatmap showing the number of cells detected in defined neuroanatomical labels.

**Figure S4.**
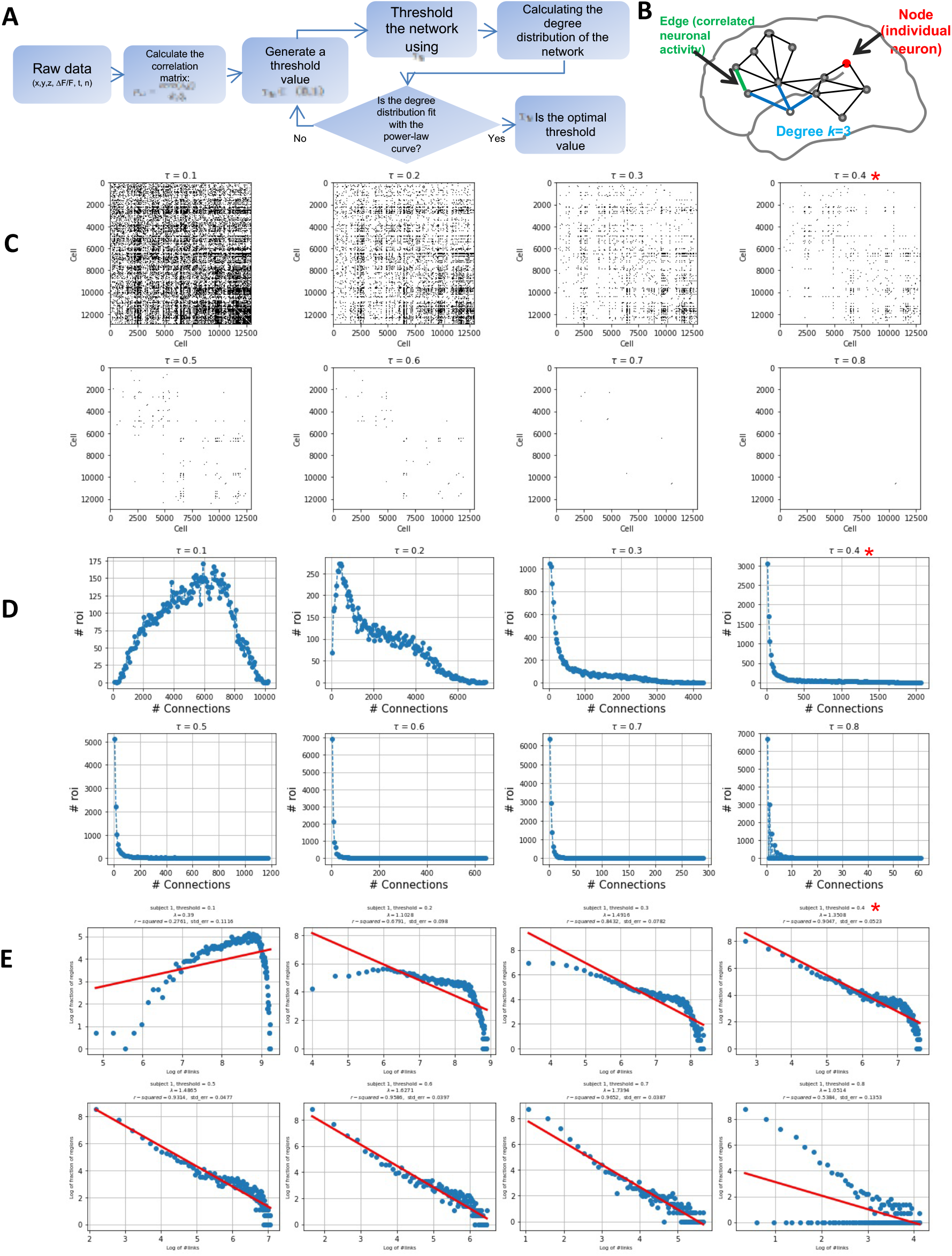
Identification of optimal threshold values to uncover significant functional connections in larval zebrafish calcium imaging data. **A,** a schematic showing the workflow of calculating functional connectivity for each ROI. **B,** a schematic of complex brain network, in which individual neurons are considered as nodes and the statistically significant relationships between each pair of nodes are known as edges. The number of edges each node has is called the degree. **C-E,** graphs for an example subject. (**C**) correlation matrices of different sparsity using different threshold values as indicated. Connections below the thresholding values are removed. **D**, Graphs showing degree distributions calculated from connectivity matrices as shown in c. (**E)** Graphs showing the approximation of line on the log-log scale to find the optimal threshold value at which data follow a power law. The optimal threshold value is 0.4 with a highest *r*^2^ (0.97)(marked with a red asterisk).

**Figure S5.**
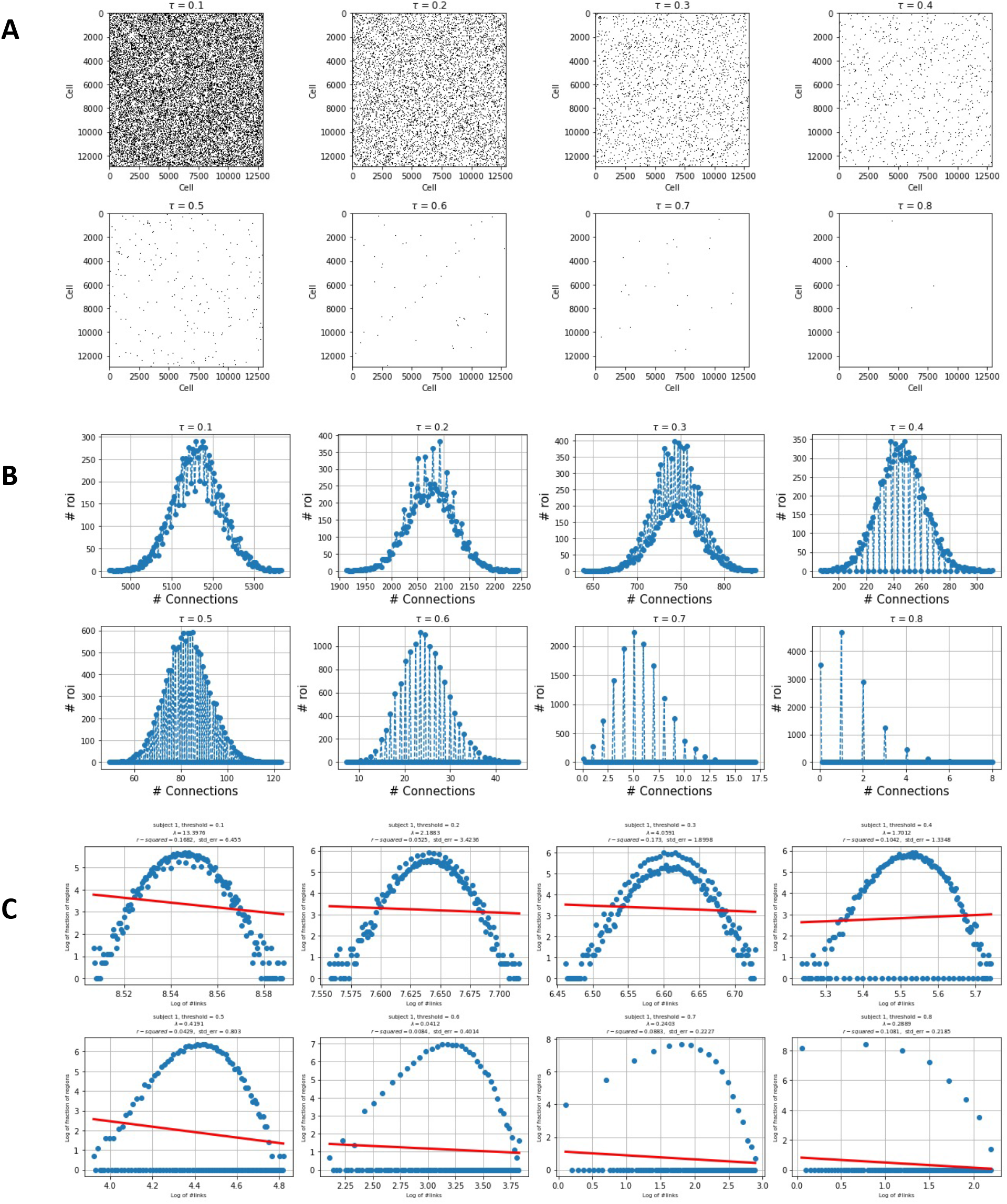
Analysis of the shuffled larval zebrafish calcium imaging data does not show power law distribution at any thresholding values. **A**, correlation matrices of different sparsity using different threshold values as indicated. Connections below the thresholding values are removed. **B**, Graphs showing degree distributions calculated from connectivity matrices as shown in A. **C**, graphs showing the approximation of line on the log-log scale. No power law distribution is observed at any thresholding value.

**Figure S6.**
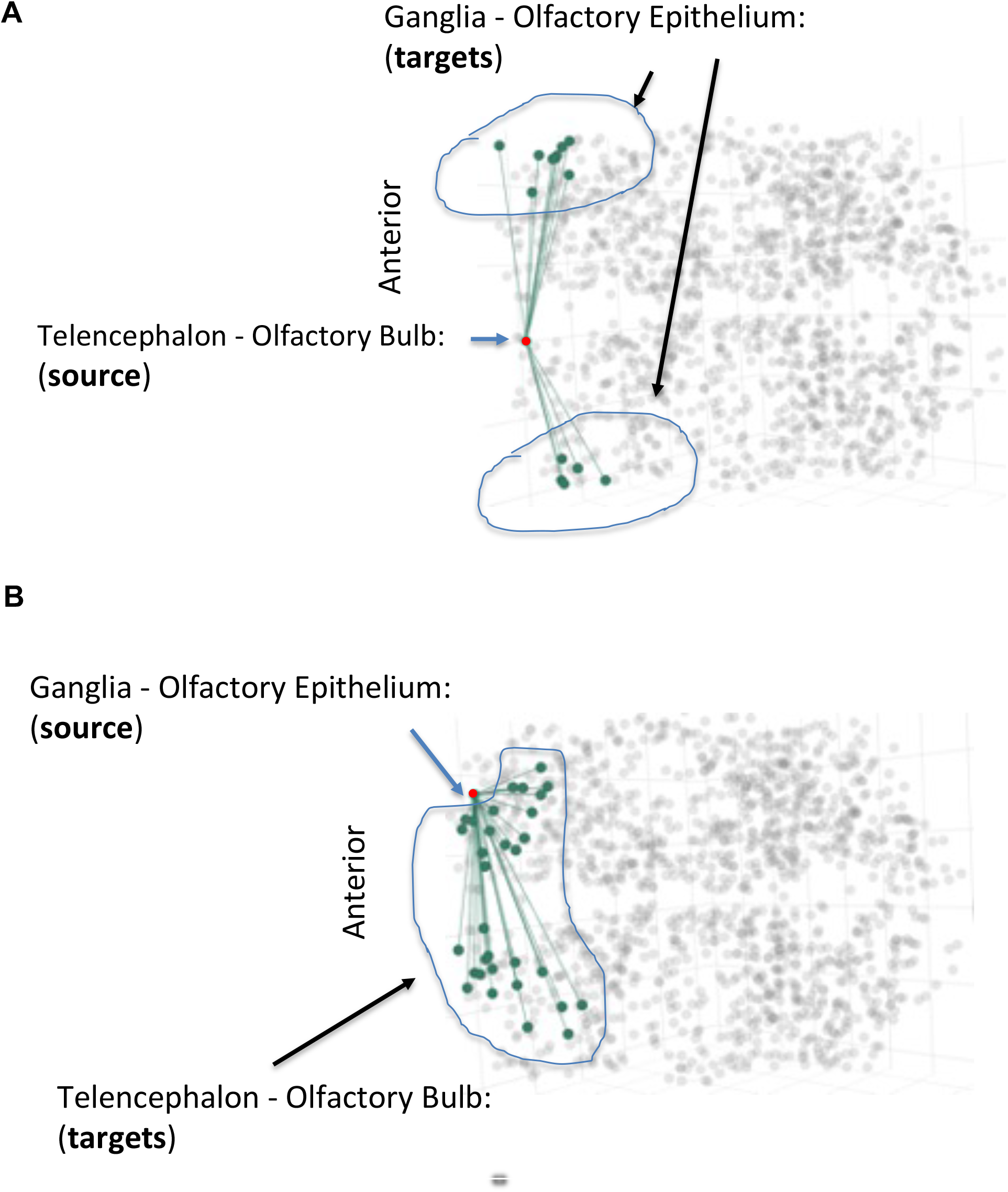
Validation of detected functional connections in the larval zebrafish forebrain. **A**, a source cell in the Telencephalon Olfactory blub is detected to have functional connections with cells in the olfactory epithelium. **B**, a source cell in the Ganglia - Olfactory Epithelium is detected to make functional connections with cells in the Telencephalon Olfactory blub. Olfactory epithelium is known to be connected with Olfactory bulb, thus validating our method of detecting functional connectivity.

**Figure S7.**
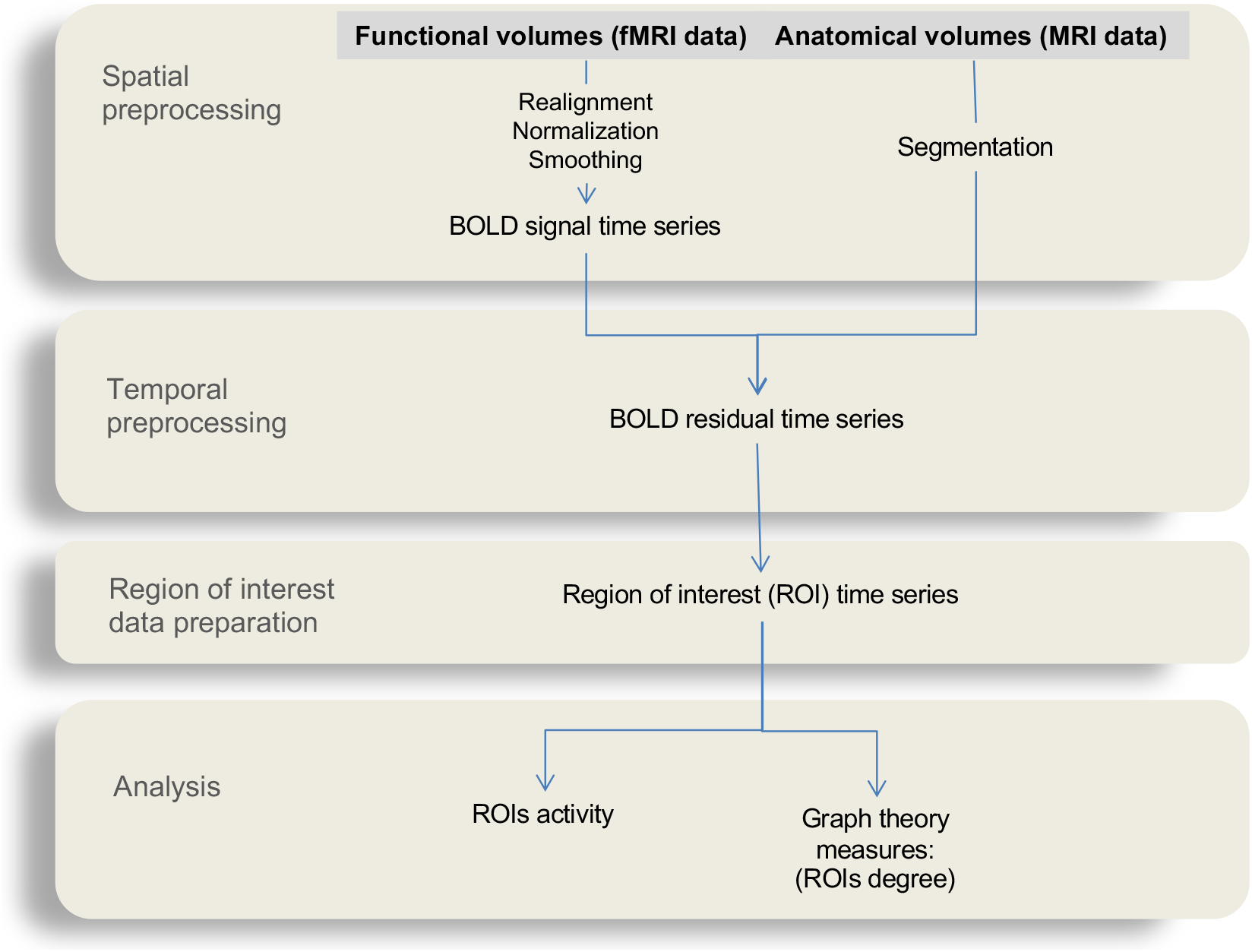
A flowchart showing the human brain data preprocessing and analysis pipeline.

**Figure S8.**
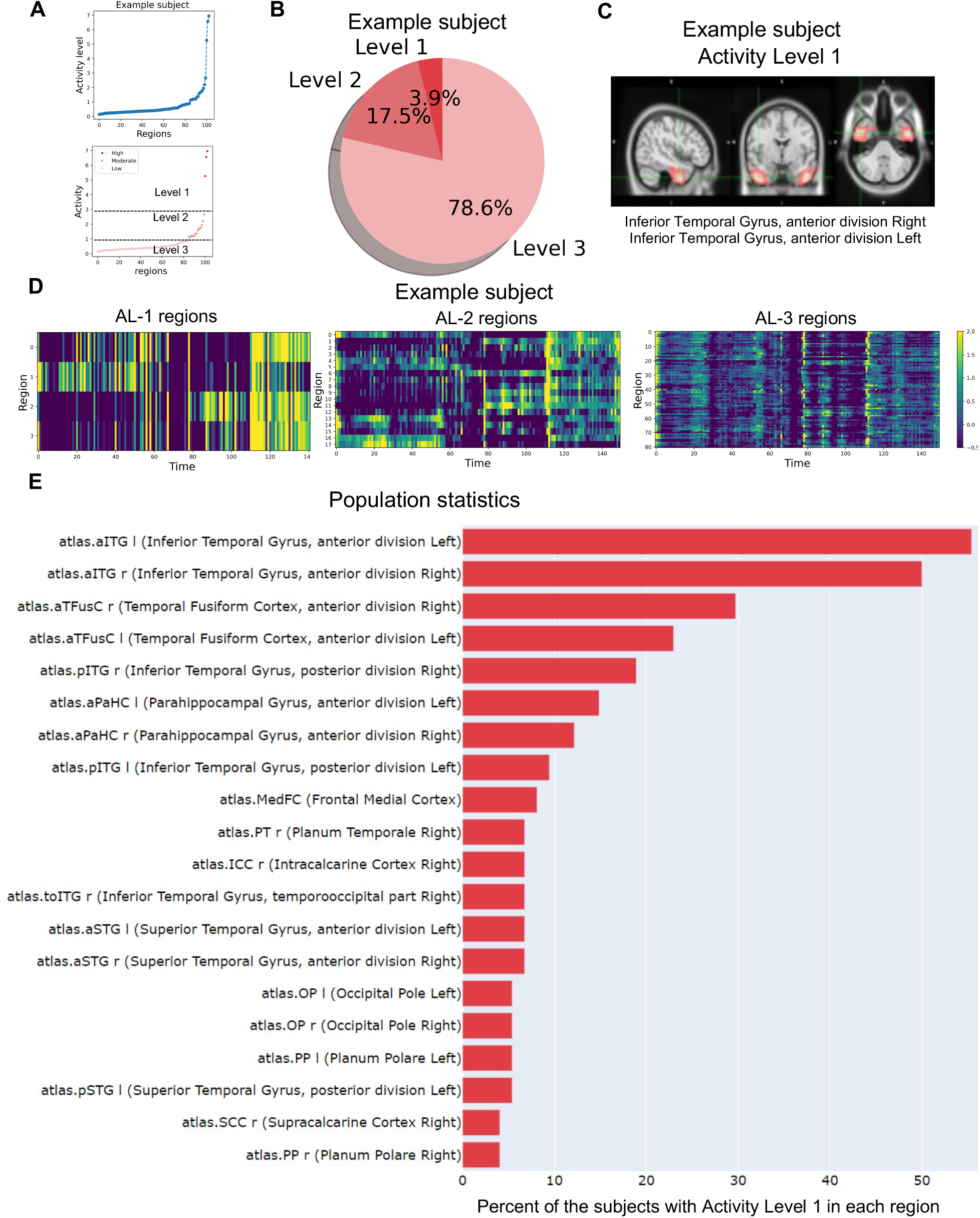
Characterization of activity in the human brain. **A,** the sorted (top) and the clustered activity levels using k-means algorithm (bottom). **B**, percentage of brain regions belonging to each activity category for an example subject. **C,** highly active brain regions (e.g., Inferior Temporal Gyrus anterior division) in an example subject, which is shared across more than 45% of the subjects. **D,** the heatmap of the Activity Level 1 (left) and Activity Level 3 brain regions (right). **E,** Bar graph showing the top 20 brain regions in Activity Level 1 and percent of subjects classified to have Level 1 activity for each brain region.

**Fig S9.**
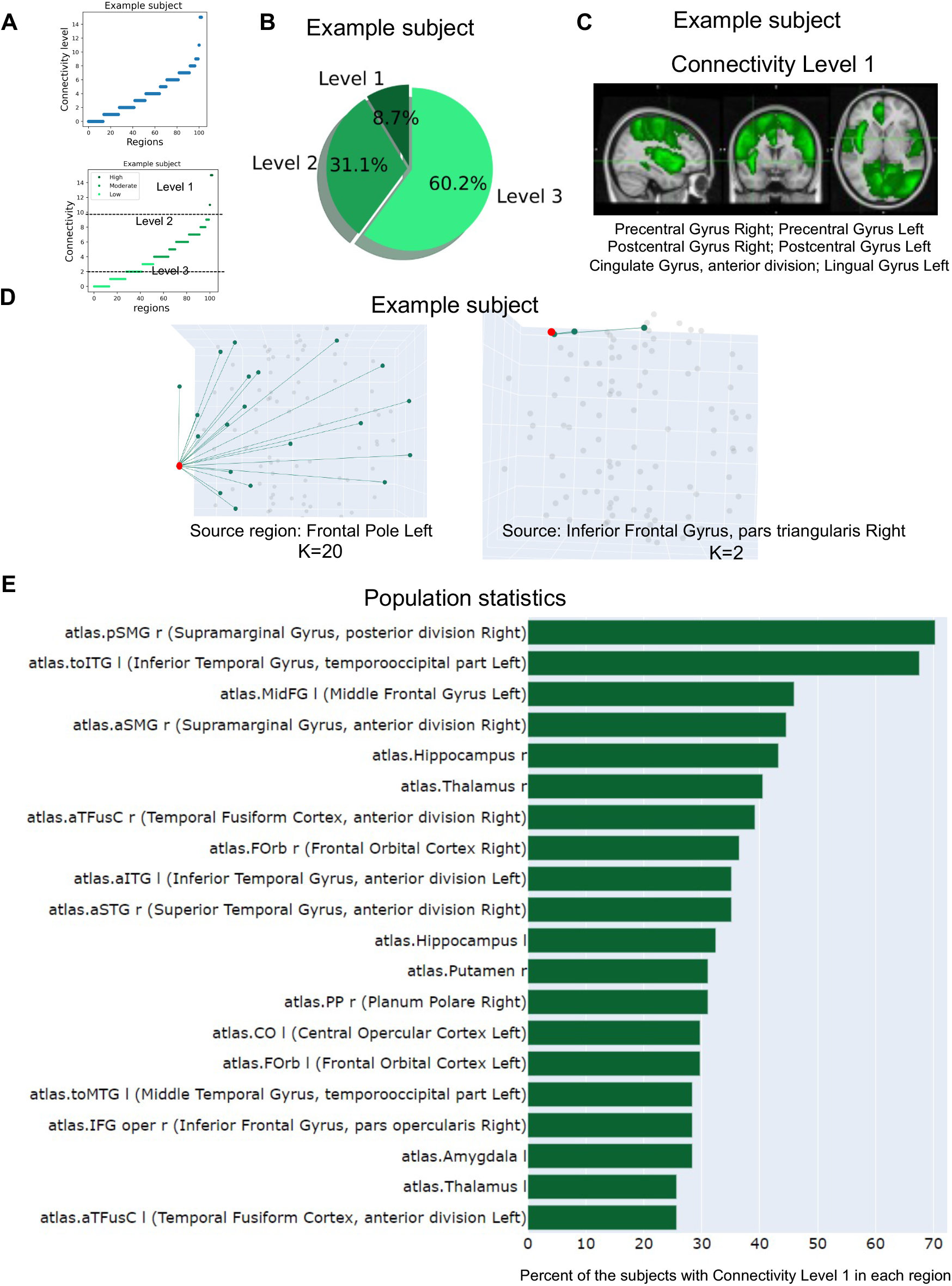
Characterization of the connectivity in the human brain. **A,** the sorted (top) and the clustered connectivity levels using k-means algorithm (bottom). **B,** percentage of brain regions in each connectivity category for an example subject. **C,** highly connected brain regions that are shared across more than 45% of the subjects. **D,** the connectivity of two example brain regions in the Connectivity Level 1 (left) and 3 (right) categories of an example subject. **E,** Percent of the subjects for each brain region with the connectivity level 1.

## References

Ahrens, M.B., Li, J.M., Orger, M.B., Robson, D.N., Schier, A.F., Engert, F., and Portugues, R. (2012). Brain-wide neuronal dynamics during motor adaptation in zebrafish. Nature 485, 471–477.

Ahrens, M.B., Orger, M.B., Robson, D.N., Li, J.M., and Keller, P.J. (2013). Whole-brain functional imaging at cellular resolution using light-sheet microscopy. Nat Methods 10, 413–420.

Avitan, L., Pujic, Z., Molter, J., Van De Poll, M., Sun, B., Teng, H., Amor, R., Scott, E.K., and Goodhill, G.J. (2017). The emergence of the spatial structure of tectal spontaneous activity Is independent of visual inputs. Curr Biol 2407–19.

Barabasi, A.L., and Albert, R. (1999). Emergence of scaling in random networks. Science 286, 509–512.

Bentley, B., Branicky, R., Barnes, C.L., Chew, Y.L., Yemini, E., Bullmore, E.T., Vértes, P.E., and Schafer, W.R. (2016). The Multilayer Connectome of Caenorhabditis elegans. PLoS Comput Biol 12, e1005283.

Biswal, B., Yetkin, F.Z., Haughton, V.M., and Hyde, J.S. (1995). Functional connectivity in the motor cortex of resting human brain using echo-planar MRI. Magn Reson Med 34, 537–541.

Briggman, K.L., Helmstaedter, M., and Denk, W. (2011). Wiring specificity in the direction-selectivity circuit of the retina. Nature 471, 183–188.

Bullmore, E., and Sporns, O. (2009). Complex brain networks: Graph theoretical analysis of structural and functional systems. Nat Rev Neurosci 10, 186–198.

Calhoun, V.D., Sui, J., Kiehl, K., Turner, J., Allen, E., and Pearlson, G. (2012). Exploring the psychosis functional connectome: aberrant intrinsic networks in schizophrenia and bipolar disorder. Front Psychiatry 2, 75.

Chen, B.C., Legant, W.R., Wang, K., Shao, L., Milkie, D.E., Davidson, M.W., Janetopoulos, C., Wu, X.S., Hammer, J.A., Liu, Z., et al. (2014). Lattice light-sheet microscopy: imaging molecules to embryos at high spatiotemporal resolution. Science 346, 1257998.

Chen, T.W., Wardill, T.J., Sun, Y., Pulver, S.R., Renninger, S.L., Baohan, A., Schreiter, E.R., Kerr, R.A., Orger, M.B., Jayaraman, V., et al. (2013). Ultrasensitive fluorescent proteins for imaging neuronal activity. Nature 499, 295–300.

Chen, X., Mu, Y., Hu, Y., Kuan, A.T., Nikitchenko, M., Randlett, O., Chen, A.B., Gavornik, J.P., Sompolinsky, H., Engert, F., et al. (2018). Brain-wide Organization of Neuronal Activity and Convergent Sensorimotor Transformations in Larval Zebrafish. Neuron 100, 876–890.

Chuang, K.H., and Nasrallah, F.A. (2017). Functional networks and network perturbations in rodents. Neuroimage 163, 419–436.

Davison, A.C., and Hinkley, D., V. (1997). Bootstrap methods and their application. (Cambridge University Press).

Deco, G., Jirsa, V.K., and McIntosh, A.R. (2011). Emerging concepts for the dynamical organization of resting-state activity in the brain. . Nat Rev Neurosci 12, 43–56.

Dunn, T.W., Gebhardt, C., Naumann, E.A., Riegler, C., Ahrens, M.B., Engert, F., and Del Bene, F. (2016). Neural Circuits Underlying Visually Evoked Escapes in Larval Zebrafish. Neuron 89, 613–628.

Fox, M.D., and Raichle, M.E. (2007). Spontaneous fluctuations in brain activity observed with functional magnetic resonance imaging. Nat Rev Neurosci 8, 700–711.

Gaudet, I., Hüsser, A., Vannasing, P., and Gallagher, A. (2020). Functional Brain Connectivity of Language Functions in Children Revealed by EEG and MEG: A Systematic Review. Front Hum Neurosci 14, 62.

Greicius, M. (2008). Resting-state functional connectivity in neuropsychiatric disorders. Curr Opin Neurol 21, 424–430.

Hagmann, P., Cammoun, L., Gigandet, X., Meuli, R., Honey, C.J., Wedeen, V.J., and Sporns, O. (2008). Mapping the structural core of human cerebral cortex. PLoS Biol 6, e159.

Hebb, D.O. (1949). The Organization of Behavior. (New York: Wiley & Sons).

Hildebrand, D.G.C., Cicconet, M., Torres, R.M., Choi, W., Quan, T.M., Moon, J., Wetzel, A.W., Scott, C.A., Graham, B.J., and al, e. (2017). Whole-brain serial-section electron microscopy in larval zebrafish. Nature 545, 345–349.

Honey, C.J., Sporns, O., Cammoun, L., Gigandet, X., Thiran, J.P., Meuli, R., and Hagmann, P. (2009). Predicting human resting-state functional connectivity from structural connectivity. Proc Natl Acad Sci U S A 106, 2035–2040.

Huisken, J., Swoger, J., Del Bene, F., Wittbrodt, J., and Stelzer, E.H. (2004). Optical sectioning deep inside live embryos by selective plane illumination microscopy. Science 305, 1007–1009.

Jernigan, T.L., and Stiles, J. (2017). Construction of the human forebrain. Wiley Interdiscip Rev Cogn Sci 8, 1–2.

Jia, H., Rochefort, N.L., Chen, X., and Konnerth, A. (2011). In vivo two-photon imaging of sensory-evoked dendritic calcium signals in cortical neurons. Nat Protoc 6, 28–35.

Kasthuri, N., Hayworth, K.J., Berger, D.R., Schalek, R.L., Conchello, J.A., Knowles-Barley, S., Lee, D., Vázquez-Reina, A., Kaynig, V., and al, e. (2015). Saturated Reconstruction of a Volume of Neocortex. Cell 162, 648–661.

Kawashima, T., Zwart, M.F., Yang, C.T., Mensh, B.D., and Ahrens, M.B. (2016). The Serotonergic System Tracks the Outcomes of Actions to Mediate Short-Term Motor Learning. Cell 167, 933–946.

Keller, P.J., Schmidt, A.D., Wittbrodt, J., and Stelzer, E.H. (2008). Reconstruction of zebrafish early embryonic development by scanned light sheet microscopy. Science 322, 1065–1069.

Lancaster, J.L., Woldorff, M.G., Parsons, L.M., Liotti, M., Freitas, C.S., Rainey, L., Kochunov, P.V., Nickerson, D., Mikiten, S.A., and Fox, P.T. (2000). Automated Talairach atlas labels for functional brain mapping. Hum Brain Mapp 10, 120–131.

Lin, Q., Manley, J., Helmreich, M., Schlumm, F., Li, J.M., Robson, D.N., Engert, F., Schier, A., Nöbauer, T., and Vaziri, A. (2020). Cerebellar Neurodynamics Predict Decision Timing and Outcome on the Single-Trial Level. Cell 180, 536–551.

MacQueen, J.B. (1967). Some Methods for classification and Analysis of Multivariate Observations. Proceedings of 5-th Berkeley Symposium on Mathematical Statistics and Probability 1, 281–297.

Meinertzhagen, I.A. (2016). Connectome studies on Drosophila: a short perspective on a tiny brain. J Neurogenet 30, 62–68.

Naumann, E.A., Fitzgerald, J.E., Dunn, T.W., Rihel, J., Sompolinsky, H., and Engert, F. (2016). From Whole-Brain Data to Functional Circuit Models: The Zebrafish Optomotor Response. Cell 167, 947–960.

Oh, S.W., Harris, J.A., Ng, L., Winslow, B., Cain, N., Mihalas, S., Wang, Q., Lau, C., Kuan, L., Henry, A.M., et al. (2014). A mesoscale connectome of the mouse brain. Nature 508, 207–214.

Pietri, T., Romano, S.A., Perez-Schuster, V., Boulanger-Weill, J., Candat, V., and Sumbre, G. (2017). The emergence of the spatial structure of tectal spontaneous activity is independent of visual inputs. Cell Reports 19, 939–948.

Ponce-Alvarez, A., Jouary, A., Privat, M., Deco, G., and Sumbre, G. (2018). Whole-Brain Neuronal Activity Displays Crackling Noise Dynamics. Neuron 100, 1–14.

Portugues, R., Feierstein, C.E., Engert, F., and Orger, M.B. (2014). Whole-brain activity maps reveal stereotyped, distributed networks for visuomotor behavior. Neuron 81, 1328–1343.

Puelles, L., and Rubenstein, J.L. (2003). Forebrain gene expression domains and the evolving prosomeric model. Trends Neurosci 26, 469–476.

Raichle, M.E., MacLeod, A.M., Snyder, A.Z., Powers, W.J., Gusnard, D.A., and Shulman, G.L. (2001). A default mode of brain function. Proc Natl Acad Sci U S A 98, 676–682.

Randlett, O., Wee, C.L., Naumann, E.A., Nnaemeka, O., Schoppik, D., Fitzgerald, J.E., Portugues, R., Lacoste, A.M., Riegler, C., Engert, F., et al. (2015). Whole-brain activity mapping onto a zebrafish brain atlas. Nat Methods 12, 1039–1046.

Rohlfing, T., and Maurer, C.R. (2003). Nonrigid image registration in shared-memory multiprocessor environments with application to brains, breasts, and bees. IEEE Trans Inform Technol Biomed 7, 16–25.

Romano, S.A., Pietri, T., Pérez-Schuster, V., Jouary, A., Haudrechy, M., and Sumbre, G. (2015). Spontaneous neuronal network dynamics reveal circuit’s functional adaptations for behavior. Neuron 85, 1070–1085.

Rubinov, M., and Sporns, O. (2010). Complex network measures of brain connectivity: Uses and interpretations. Neuroimage 52, 1059–1069.

Shulman, R.G., Rothman, D.L., Behar, K.L., and Hyder, F. (2004). Energetic basis of brain activity: implications for neuroimaging. Trends Neurosci 27, 489–495.

Sporns, O., and Betzel, R.F. (2016). Modular Brain Networks. Annu Rev Psychol 67, 613–640.

Tosches, M.A., and Arendt, D. (2013). The bilaterian forebrain: an evolutionary chimaera. Curr Opin Neurobiol 23, 1080–1089.

Vanwalleghem, G.C., Ahrens, M.B., and Scott, E.K. (2018). Integrative whole-brain neuroscience in larval zebrafish. Curr Opin Neurobiol 50, 136–145.

Wanner, A.A., Genoud, C., Masudi, T., Siksou, L., and Friedrich, R.W. (2016). Dense EM-based reconstruction of the interglomerular projectome in the zebrafish olfactory bulb. Nat Neurosci 19, 816–825.

Watts, D.J., and Strogatz, S.H. (1998). Collective dynamics of ‘small-world’ networks. Nature 393, 440.

White, J.G., Southgate, E., Thomson, J.N., and Brenner, S. (1986). The structure of the nervous system of the nematode Caenorhabditis elegans. . Philos Trans R Soc Lond B314, 1–340.

Wilson, S.W., and Houart, C. (2004). Early steps in the development of the forebrain. Dev Cell 6, 167–181.

Wu, Y., Ghitani, A., Christensen, R., Santella, A., Du, Z., Rondeau, G., Bao, Z., Colón-Ramos, D., and Shroff, H. (2011). Inverted selective plane illumination microscopy (iSPIM) enables coupled cell identity lineaging and neurodevelopmental imaging in Caenorhabditis elegans. Proc Natl Acad Sci 108, 17708–17713.

Zingg, B., Hintiryan, H., Gou, L., Song, M.Y., Bay, M., Bienkowski, M.S., Foster, N.N., Yamashita, S., Bowman, I., Toga, A.W., et al. (2014). Neural networks of the mouse neocortex. Cell 156, 1096–1111.

## References cited

1. J. G. White, E. Southgate, J. N. Thomson, S. Brenner, The structure of the nervous system of the nematode Caenorhabditis elegans. . Philos. Trans. R. Soc. Lond. B314, 1–340 (1986).

2. I. A. Meinertzhagen, Connectome studies on Drosophila: a short perspective on a tiny brain. J. Neurogenet. 30, 62–68 (2016).

3. G. C. Vanwalleghem, M. B. Ahrens, E. K. Scott, Integrative whole-brain neuroscience in larval zebrafish. Curr. Opin. Neurobiol. 50, 136–145 (2018).

4. K. H. Chuang, F. A. Nasrallah, Functional networks and network perturbations in rodents. Neuroimage 163, 419–436 (2017).

5. O. Sporns, R. F. Betzel, Modular Brain Networks. Annu. Rev. Psychol. 67, 613–640 (2016).

6. M. Rubinov, O. Sporns, Complex network measures of brain connectivity: Uses and interpretations. Neuroimage 52, 1059–1069 (2010).

7. E. Bullmore, O. Sporns, Complex brain networks: Graph theoretical analysis of structural and functional systems. Nat. Rev. Neurosci. 10, 186–198 (2009).

8. S. W. Oh et al., A mesoscale connectome of the mouse brain. Nature 508, 207–214 (2014).

9. B. Zingg et al., Neural networks of the mouse neocortex. Cell 156, 1096–1111 (2014).

10. P. Hagmann et al., Mapping the structural core of human cerebral cortex. PLoS Biol. 6, e159 (2008).

11. D. G. C. Hildebrand et al., Whole-brain serial-section electron microscopy in larval zebrafish. Nature 545, 345–349 (2017).

12. N. Kasthuri et al., Saturated Reconstruction of a Volume of Neocortex. Cell 162, 648–661 (2015).

13. A. A. Wanner, C. Genoud, T. Masudi, L. Siksou, R. W. Friedrich, Dense EM-based reconstruction of the interglomerular projectome in the zebrafish olfactory bulb. Nat. Neurosci. 19, 816–825 (2016).

14. K. L. Briggman, M. Helmstaedter, W. Denk, Wiring specificity in the direction-selectivity circuit of the retina. Nature 471, 183–188 (2011).

15. D. O. Hebb, The Organization of Behavior., (Wiley & Sons, New York, 1949).

16. C. J. Honey et al., Predicting human resting-state functional connectivity from structural connectivity. Proc. Natl. Acad. Sci. U S A. 106, 2035–2040 (2009).

17. B. Bentley et al., The Multilayer Connectome of Caenorhabditis elegans. PLoS Comput. Biol. 12, e1005283 (2016).

18. I. Gaudet, A. Hüsser, P. Vannasing, A. Gallagher, Functional Brain Connectivity of Language Functions in Children Revealed by EEG and MEG: A Systematic Review. Front. Hum. Neurosci. 14, 62 (2020).

19. T. W. Chen et al., Ultrasensitive fluorescent proteins for imaging neuronal activity. Nature 499, 295–300 (2013).

20. J. Huisken, J. Swoger, F. Del Bene, J. Wittbrodt, E. H. Stelzer, Optical sectioning deep inside live embryos by selective plane illumination microscopy. Science 305, 1007–1009 (2004).

21. P. J. Keller, A. D. Schmidt, J. Wittbrodt, E. H. Stelzer, Reconstruction of zebrafish early embryonic development by scanned light sheet microscopy. Science 322, 1065–1069 (2008).

22. B. C. Chen et al., Lattice light-sheet microscopy: imaging molecules to embryos at high spatiotemporal resolution. Science 346, 1257998 (2014).

23. M. B. Ahrens et al., Brain-wide neuronal dynamics during motor adaptation in zebrafish. Nature 485, 471–477 (2012).

24. M. B. Ahrens, M. B. Orger, D. N. Robson, J. M. Li, P. J. Keller, Whole-brain functional imaging at cellular resolution using light-sheet microscopy. Nat. Methods 10, 413–420 (2013).

25. R. Portugues, C. E. Feierstein, F. Engert, M. B. Orger, Whole-brain activity maps reveal stereotyped, distributed networks for visuomotor behavior. Neuron 81, 1328–1343 (2014).

26. E. A. Naumann et al., From Whole-Brain Data to Functional Circuit Models: The Zebrafish Optomotor Response. Cell 167, 947–960 (2016).

27. X. Chen et al., Brain-wide Organization of Neuronal Activity and Convergent Sensorimotor Transformations in Larval Zebrafish. Neuron 100, 876–890 (2018).

28. T. W. Dunn et al., Neural Circuits Underlying Visually Evoked Escapes in Larval Zebrafish. Neuron 89, 613–628 (2016).

29. T. Kawashima, M. F. Zwart, C. T. Yang, B. D. Mensh, M. B. Ahrens, The Serotonergic System Tracks the Outcomes of Actions to Mediate Short-Term Motor Learning. Cell 167, 933–946 (2016).

30. Q. Lin et al., Cerebellar Neurodynamics Predict Decision Timing and Outcome on the Single-Trial Level. Cell 180, 536–551 (2020).

31. S. A. Romano et al., Spontaneous neuronal network dynamics reveal circuit’s functional adaptations for behavior. Neuron 85, 1070–1085 (2015).

32. A. Ponce-Alvarez, A. Jouary, M. Privat, G. Deco, G. Sumbre, Whole-Brain Neuronal Activity Displays Crackling Noise Dynamics. Neuron 100, 1–14 (2018).

33. L. Avitan et al., The emergence of the spatial structure of tectal spontaneous activity Is independent of visual inputs. Curr. Biol. 2407–19, (2017).

34. T. Pietri et al., The emergence of the spatial structure of tectal spontaneous activity is independent of visual inputs. Cell Reports 19, 939–948 (2017).

35. G. Deco, V. K. Jirsa, A. R. McIntosh, Emerging concepts for the dynamical organization of resting-state activity in the brain. . Nat. Rev. Neurosci. 12, 43–56 (2011).

36. B. Biswal, F. Z. Yetkin, V. M. Haughton, J. S. Hyde, Functional connectivity in the motor cortex of resting human brain using echo-planar MRI. Magn. Reson. Med. 34, 537–541 (1995).

37. M. E. Raichle et al., A default mode of brain function. Proc. Natl. Acad. Sci. U S A 98, 676–682 (2001).

38. M. Greicius, Resting-state functional connectivity in neuropsychiatric disorders. Curr. Opin. Neurol. 21, 424–430 (2008).

39. Y. Wu et al., Inverted selective plane illumination microscopy (iSPIM) enables coupled cell identity lineaging and neurodevelopmental imaging in Caenorhabditis elegans. Proc. Natl. Acad. Sci. 108, 17708–17713 (2011).

40. S. W. Wilson, C. Houart, Early steps in the development of the forebrain. Dev. Cell 6, 167–181 (2004).

41. L. Puelles, J. L. Rubenstein, Forebrain gene expression domains and the evolving prosomeric model. Trends Neurosci. 26, 469–476 (2003).

42. T. L. Jernigan, J. Stiles, Construction of the human forebrain. Wiley Interdiscip. Rev. Cogn. Sci. 8, 1–2 (2017).

43. M. A. Tosches, D. Arendt, The bilaterian forebrain: an evolutionary chimaera. Curr. Opin. Neurobiol. 23, 1080–1089 (2013).

44. O. Randlett et al., Whole-brain activity mapping onto a zebrafish brain atlas. Nat. Methods 12, 1039–1046 (2015).

45. H. Jia, N. L. Rochefort, X. Chen, A. Konnerth, In vivo two-photon imaging of sensory-evoked dendritic calcium signals in cortical neurons. Nat. Protoc. 6, 28–35 (2011).

46. T. Rohlfing, C. R. Maurer, Nonrigid image registration in shared-memory multiprocessor environments with application to brains, breasts, and bees. IEEE Trans. Inform. Technol. Biomed. 7, 16–25 (2003).

47. J. B. MacQueen, Some Methods for classification and Analysis of Multivariate Observations. Proceedings of 5-th Berkeley Symposium on Mathematical Statistics and Probability 1, 281–297 (1967).

48. D. J. Watts, S. H. Strogatz, Collective dynamics of ‘small-world’ networks. Nature 393, 440 (1998).

49. A. L. Barabasi, R. Albert, Emergence of scaling in random networks. Science 286, 509–512 (1999).

50. A. C. Davison, D. Hinkley, v., Bootstrap methods and their application., (Cambridge University Press 1997).

51. V. D. Calhoun et al., Exploring the psychosis functional connectome: aberrant intrinsic networks in schizophrenia and bipolar disorder. Front. Psychiatry 2, 75 (2012).

52. J. L. Lancaster et al., Automated Talairach atlas labels for functional brain mapping. Hum. Brain Mapp. 10, 120–131 (2000).

53. M. D. Fox, M. E. Raichle, Spontaneous fluctuations in brain activity observed with functional magnetic resonance imaging. Nat. Rev. Neurosci. 8, 700–711 (2007).

54. R. G. Shulman, D. L. Rothman, K. L. Behar, F. Hyder, Energetic basis of brain activity: implications for neuroimaging. Trends Neurosci. 27, 489–495 (2004).

55. A. Kumar et al., Dual-view plane illumination microscopy for rapid and spatially isotropic imaging. . Nat. Protoc. 9, 2555–2573 (2014).

56. P. Thévenaz, U. E. Ruttimann, M. Unser, A pyramid approach to subpixel registration based on intensity. IEEE Transactions on Image Processing 7, 27–41 (1998).

57. S. van der Walt et al., Scikit-image: image processing in Python. Peer J. 2, e453 (2014).

58. N. W. Churchill et al., Optimizing preprocessing and analysis pipelines for single-subject fMRI. I. Standard temporal motion and physiological noise correction methods. Hum. Brain Mapp. 33, 609–627 (2012).

59. Y. Behzadi, K. Restom, J. Liau, T. T. Liu, A component based noise correction method (CompCor) for BOLD and perfusion based fMRI. Neuroimage 37, 90–101 (2007).

60. W. S. CLeveland, Robust locally weighted regression and smoothing scatterplo. Journal of the American Statistical Association 74, 829–836 (1979).

## References cited

1. Y. Wu et al., Inverted selective plane illumination microscopy (iSPIM) enables coupled cell identity lineaging and neurodevelopmental imaging in Caenorhabditis elegans. Proc. Natl. Acad. Sci. 108, 17708–17713 (2011).

2. A. Kumar et al., Dual-view plane illumination microscopy for rapid and spatially isotropic imaging. . Nat. Protoc. 9, 2555–2573 (2014).

3. P. Thévenaz, U. E. Ruttimann, M. Unser, A pyramid approach to subpixel registration based on intensity. IEEE Transactions on Image Processing 7, 27–41 (1998).

4. S. van der Walt et al., Scikit-image: image processing in Python. Peer J. 2, e453 (2014).

5. H. Jia, N. L. Rochefort, X. Chen, A. Konnerth, In vivo two-photon imaging of sensory-evoked dendritic calcium signals in cortical neurons. Nat. Protoc. 6, 28–35 (2011).

6. S. A. Romano et al., Spontaneous neuronal network dynamics reveal circuit’s functional adaptations for behavior. Neuron 85, 1070–1085 (2015).

7. O. Randlett et al., Whole-brain activity mapping onto a zebrafish brain atlas. Nat. Methods 12, 1039–1046 (2015).

8. N. W. Churchill et al., Optimizing preprocessing and analysis pipelines for single-subject fMRI. I. Standard temporal motion and physiological noise correction methods. Hum. Brain Mapp. 33, 609–627 (2012).

9. Y. Behzadi, K. Restom, J. Liau, T. T. Liu, A component based noise correction method (CompCor) for BOLD and perfusion based fMRI. Neuroimage 37, 90–101 (2007).

10. J. L. Lancaster et al., Automated Talairach atlas labels for functional brain mapping. Hum. Brain Mapp. 10, 120–131 (2000).

11. J. B. MacQueen, Some Methods for classification and Analysis of Multivariate Observations. Proceedings of 5-th Berkeley Symposium on Mathematical Statistics and Probability 1, 281–297 (1967).

12. M. Rubinov, O. Sporns, Complex network measures of brain connectivity: Uses and interpretations. Neuroimage 52, 1059–1069 (2010).

13. W. S. CLeveland, Robust locally weighted regression and smoothing scatterplo. Journal of the American Statistical Association 74, 829–836 (1979).

14. A. C. Davison, D. Hinkley, V., Bootstrap methods and their application., (Cambridge University Press 1997).

